# *Cascabel*: a flexible, scalable and easy-to-use amplicon sequence data analysis pipeline

**DOI:** 10.1101/809384

**Authors:** Alejandro Abdala Asbun, Marc A Besseling, Sergio Balzano, Judith van Bleijswijk, Harry Witte, Laura Villanueva, Julia C Engelmann

## Abstract

Marker gene sequencing of the rRNA operon (16S, 18S, ITS) or cytochrome c oxidase I (CO1) is a popular means to assess microbial communities of the environment, microbiomes associated with plants and animals, as well as communities of multicellular organisms via environmental DNA sequencing. Since this technique is based on sequencing a single gene rather than the entire genome, the number of reads needed per sample is lower than that required for metagenome sequencing, making marker gene sequencing affordable to nearly any laboratory. Despite the relative ease and cost-efficiency of data generation, analyzing the resulting sequence data requires computational skills that may go beyond the standard repertoire of a current molecular biologist/ecologist. We have developed *Cascabel*, a flexible and easy-to-use amplicon sequence data analysis pipeline, which uses Snakemake and a combination of existing and newly developed solutions for its computational steps. *Cascabel* takes the raw data as input and delivers a table of operational taxonomic units (OTUs) and a representative sequence tree. Our pipeline allows customizing the analyses by offering several choices for most of the steps, for example different OTU generating methods. The pipeline can make use of multiple computing nodes and scales from personal computers to computing servers. The analyses and results are fully reproducible and documented in an HTML and optional pdf report. *Cascabel* is freely available at Github: https://github.com/AlejandroAb/CASCABEL and licensed under GNU GPLv3.

## Introduction

High-throughput sequencing of an omnipresent marker gene such as the gene coding for the small subunit of the ribosomal RNA (16S for prokaryotes or 18S for eukaryotes) is a cheap means for microbial community profiling that is affordable for nearly every lab. Moreover, sequencing a marker gene like cytochrome c oxidase I (CO1) in environmental DNA also allows to track larger multicellular organisms, for example fish in the sea. Amplicon sequencing can also be used to investigate active microbial communities based on ribosomal RNA abundance instead of the rRNA gene locus (1, 2). Typically, a short fragment of 100-600 nucleotides of the marker gene with the desired taxonomic resolution is amplified by PCR from the DNA extract or cDNA of the community and then sequenced by high throughput sequencing. On current sequencing platforms, up to hundreds of samples can be combined (multiplexed) in a single sequencing run, decreasing the sequencing costs per sample tremendously. Not surprisingly, community compositions based on DNA analyses have been generated from most of the habitats on earth, including the human body, e.g., (3), the open ocean, e.g., (4), deep sea, e.g., (5) and intracellular symbionts, e.g., (6). The current bottleneck in studies using community profiling is the computational analysis of the (potentially massive) sequence data. For scientists with little background in bioinformatics, the amount of data and complexity of data analysis can be overwhelming. Several software solutions for the individual steps from raw sequence data to an operational taxonomic unit (OTU) table exist (7, 8), but are not necessarily straightforward to use. The software package Mothur (7) which comes with its own computational environment and the QIIME framework (8) have been popular platforms for data analysis, but both solutions require the ability to work on the command line. Analyzing multiple sequencing libraries quickly becomes tedious for users not proficient in implementing bash (or any other programming language) scripts which chain the individual steps and allow parallel processing. While web servers for microbial community data analysis like SILVAngs (9) and MG-RAST (10) are easy to use, they are inherently inflexible and also limited in throughput. QIIME2 (11) has command line and graphical user interface modes of operation and offers even a larger choice of algorithms for data analysis than the original QIIME, including statistical analyses of the resulting community profiles. We anticipated a need for a pipeline which combines the flexibility provided by using bioinformatic tools on the command line within one of the existing frameworks with the ease of using interactive web servers for analyzing and interpreting amplicon sequencing data.

We here provide *Cascabel*, a Snakemake (12) pipeline for the analysis of community marker gene sequence data which is easy to use for people with little bioinformatics background, and both flexible and powerful enough to be attractive for people with bioinformatics training. *Cascabel* supports large sample and sequencing library throughput as well as parallel computing on personal computers and computing servers. Moreover, all input and output files, tools, parameters and their versions are documented and render the analyses fully reproducible.

## Implementation

Our pipeline makes use of the workflow management engine Snakemake (12), which scales from personal workstations to compute clusters. *Cascabel* consists of a set of ‘rules’, which specify the input, the action to perform on the input (executed by a bash/python/R/java script), and the output. The user defines via a configuration file (called ‘config file’ from now on) in yaml format, how these ‘rules’ are chained to perform amplicon sequence data analysis from the raw data to the final OTU table. For most of the rules *Cascabel* provides several alternative algorithms or tools and allow passing arguments via the config file to the algorithm being used. In addition, rules can be skipped, and the pipeline can be entered and exited at every step. This makes *Cascabel* very flexible and highly customizable. Moreover, the pipeline is easily extendable and amendable to personal needs, allowing e.g. the analysis of any marker gene sequence data. Running *Cascabel* requires the raw fastq sequence data files, a mapping file indicating which sample carries which barcode if the data should be demultiplexed, and optionally sample meta-data (e.g., geographic coordinates of the sampling stations, physical, chemical, or biological properties), and the config file specifying the tools and parameters used for running the pipeline. We provide default parameters but strongly advise making informed choices about parameter settings matching the individual needs of the experiment and data set. With the files in place, *Cascabel* is started with a one-line command on the terminal. Snakemake takes care of executing the rules in a computationally efficient manner, making optimal use of available resources e.g., distributing jobs over several nodes. Figure 1 provides an overview of the workflow of *Cascabel*. *Cascabel* supports paired-end sequence data as input from one or multiple samples per input file. Barcodes for demultiplexing samples can be situated at the beginning of one or both of the reads.

**Fig. 1.**
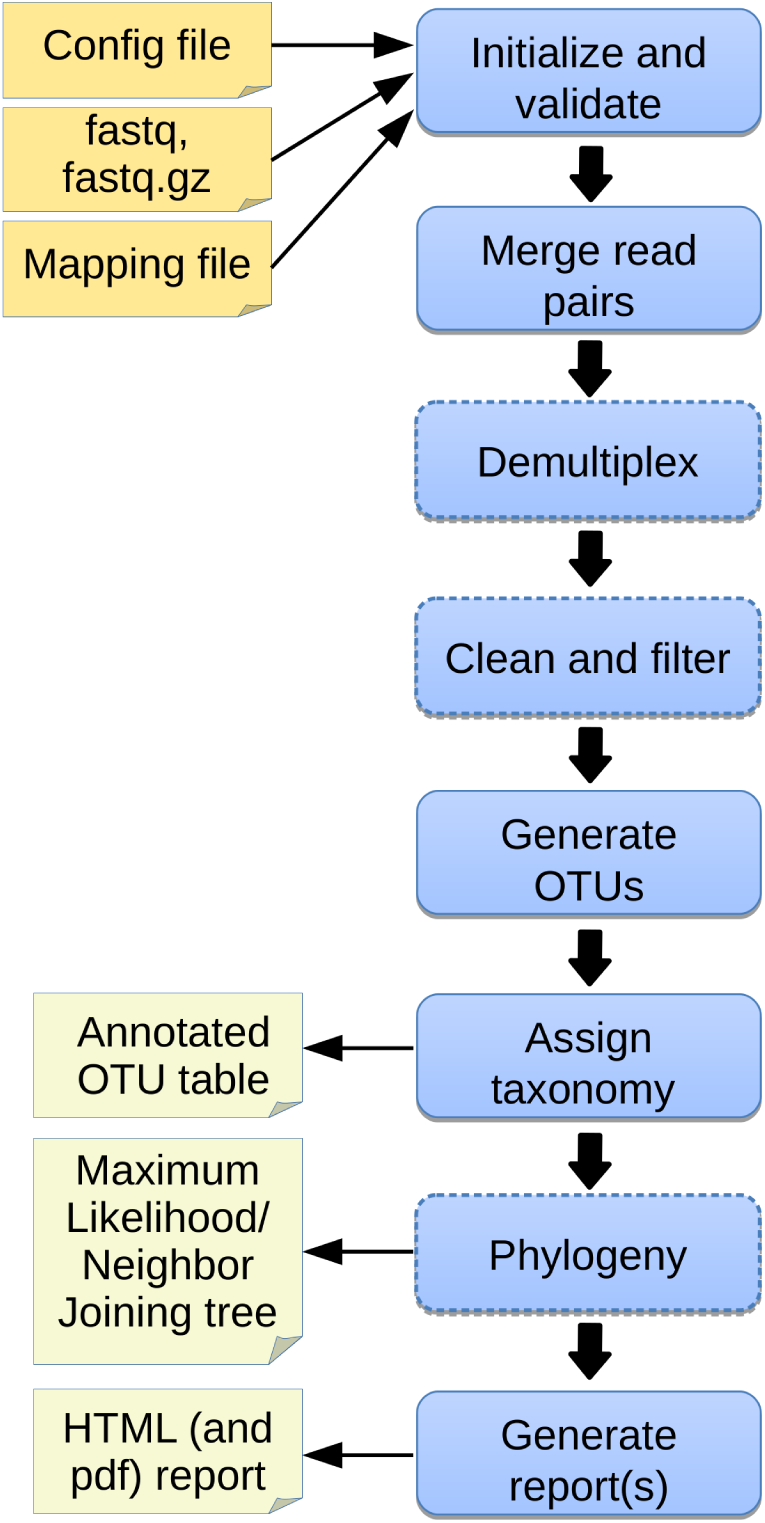
Overview of *Cascabel*. The workflow indicates input files (config file, sequence data in fastq format, barcode mapping file), mandatory and optional steps of the pipeline (blue boxes) as well as the main output files. The boxes of optional steps have dashed borders. ‘Clean and filter’ refers to removing primers/adapters and chimeras. Table 1 shows a detailed summary of the steps, available tools and output files.

*Cascabel* provides a range of popular methods to generate OTUs with or without a reference sequence database (swarm (13), sortmerna (14), mothur (7), trie (QIIME team, unpublished), uclust/uclust_ref/usearch/usearch_ref (15), prefix/suffix (QIIME team, unpublished), cd-hit (16), sumaclust (17)) which are executed by QIIME. Then, representative sequences are chosen for each OTU (with options: random, longest, most_abundant, first) (8).

**Table 1.** Outline of the steps performed by *Cascabel*. ‘script(s)’ refers to *Cascabel* scripts in bash, java or R.

| Step | Tools/Algorithms | Output |
| --- | --- | --- |
| Initialize structure | script | Project folder and file structure |
| Quality Control | FastQC(18) | FastQC report |
| Merge reads | PEAR(19) | Merged (assembled) sequences |
| Demultiplex | QIIME(8), scripts | Sequences assigned to samples in one file and per sample |
| Align versus reference | mothur(7) | Aligned sequences |
| Remove chimeras | usearch61(15) | Chimera-free sequences |
| Remove adapters | Cutadapt(20) | Adapter-free sequences |
| Size filter | script | Filtered sequences |
| Dereplicate | VSEARCH(21) | Dereplicated sequences |
| Generate OTUs | VSEARCH, swarm(13), uclust(15), trie, sortmerna(14) | OTU table |
| Assign taxonomy | QIIME (BLAST(22), uclust, RDP(23)), blastn (BLAST+)(24), VSEARCH | Taxonomic assignments for each OTU |
| Generate OTU table | QIIME | Annotated OTU table |
| Alignment | pynast(25), mafft(26), infernal(27), clustalw(28), muscle(29) | multiple sequence alignment |
| Make tree | muscle, clustalw, raxml(30), fasttree(31) | phylogenetic tree |
| Report | scripts, Krona(32) | HTML, pdf report, Krona charts |

*Cascabel* assigns taxonomy to the representative sequences using the in-built most recent version of the SILVA database, at this moment v132 (9), providing three different approaches: VSEARCH, which performs global alignment of the target sequences against the reference database; BLAST, making use of BLAST+ (24); QIIME, with methods BLAST (22), uclust or the RDP classifier. Alternatively, any other public or custom database can be used for taxonomic annotation. If taxonomy is assigned with VSEARCH or BLAST, the user can choose to assign the sequences to the lowest common ancestor (LCA) with the stampa approach (https://github.com/frederic-mahe/stampa). The last rule of *Cascabel* (the ‘target’ rule) generates HTML and optional pdf reports with documentation, figures and tables summarizing the results of individual rules, as well as all software versions used. Table 1 shows an overview of the options and methods provided for the individual steps of the analysis performed by *Cascabel*.

*Cascabel* has an interactive and a non-interactive mode. In interactive mode, several modules have a check-point which needs to be passed to continue with the analysis. If the check fails (e.g., if too many FASTQC quality modules failed or the number of sequences assigned to sample barcodes is too low), the pipeline stops and the user has to either change parameters and continue, or exit the pipeline. If parameters were changed interactively, the new ones are documented in the reports.

Snakemake will not re-run a rule if the output file of that rule already exists. This avoids over-writing existing results, but also renders it impossible to keep results of multiple analyses on the same data in the same project. To avoid initializing multiple projects with the same raw data, we implemented *Cascabel* with a ‘Run’ parameter. Whenever the user changes the Run parameter, a new analysis will be performed (except for quality control on the raw data) and the results saved in a different ‘Run’ folder. Moreover, the user can perform taxonomic assignments for the same run using different methods and the results will be saved in individual ‘taxonomy’ folders. When starting a new taxonomic assignment, the existing OTU representative sequences are used so no processing time is wasted by performing the same upstream rules several times.

## Results

To show the utility of our pipeline, we applied it to 16S rRNA gene amplicon data generated from water column samples taken from Lake Chala, which is situated on the border of Kenya and Tanzania, east of Mount Kilimanjaro in Africa. As stated earlier, however, the pipeline can process sequence data from any marker gene. *Cascabel* comes with taxonomic mapping files for 16S rRNA and 18S rRNA gene sequences from SILVA v132, but the user can always choose to make use of a different public or a custom reference sequence database.

First, *Cascabel* checked the validity of the input files including the barcode mapping file and the config file. Supplementary file 1 contains the config file of the analyses presented here. After having validated the input files, *Cascabel* proceeded with analyzing sequence data quality with FastQC (18). In interactive mode, *Cascabel* will stop if more than a specified number of quality check modules failed. Next, we assembled read pairs using PEAR (19) and assessed the quality of the assembled reads with FastQC.

When working with large datasets, a dereplication step which collapses identical sequences into one representative sequence can drastically reduce computation time. We provide a custom rule based on VSEARCH (21) which keeps track of the abundance of the individual sequence across samples. We dereplicated sequences which were identical over the full sequence length. If the library contains sequences from several samples, they are then demultiplexed based on the barcode sequences provided in the barcode mapping file. To do so, we make use of QIIME (8) and a custom R script to (optionally) allow sequence errors in the barcodes. Demultiplexed data can also be stored in individual fastq files for further use outside the pipeline, e.g., for submitting data to public repositories. We demultiplexed the example 16S rRNA gene sequencing data based on a sample barcode of 12 nucleotides located at the beginning of the forward read. Optionally, *Cascabel* will align sequence reads against a reference sequence database to facilitate removing sequence adapters or primers or both. Adapter and primer sequences can be trimmed off with Cutadapt (20). We did not apply Cutadapt on our example data set because we did not expect adapters and did not consider primer removal necessary for the purpose of demonstrating the features of *Cascabel*. Then, *Cascabel* generates a histogram of sequence lengths. In interactive mode, *Cascabel* shows the frequency of occurrence of each of the read lengths on the terminal and allows to change the minimum and maximum sequence length provided in the config file. For our 16S data set, we filtered out sequences whose length differed by more than 15 nucleotides from the average sequence length. The library report contains a smoothed histogram of the sequence lengths to validate the choice of the minimum and maximum sequence length (Figure 2A). Optionally, *Cascabel* identifies and removes chimeras either *de novo* based on sequence abundance or searching against the gold database provided by QIIME with the usearch61 algorithm (15). The user can also provide a different databases such as SILVA (9) or PRD2 (33) to search chimeras. Assembled and potentially filtered sequence reads from all samples are then concatenated into one fasta file. *Cascabel* then generates a histogram to visualize the number of reads per sample for the report of the complete run including all libraries (named ‘otu_report’) to assess whether the sequences are evenly spread across the samples (Figure 2B). In the data set analyzed, indeed, the number of reads per sample vary to some extent. This is observed frequently even though amplicons from the different samples were pooled in equimolar amounts. Furthermore, the reports for each of the libraries contain a plot of the number and percentages of raw, assembled, demultiplexed and length filtered sequences (Figure 2C).

**Fig. 2.**
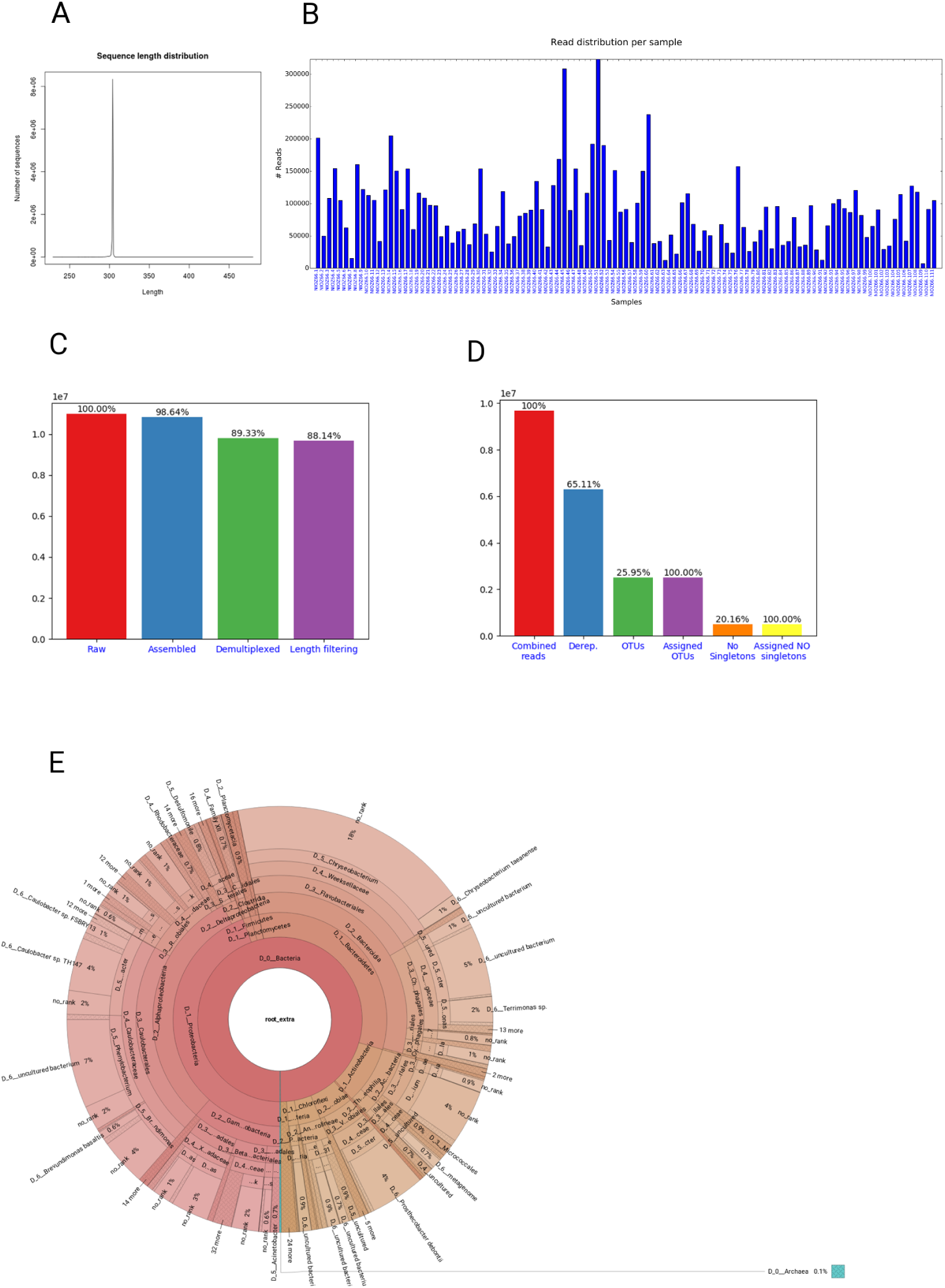
Figures shown in *Cascabel* reports. **(A)** Smoothed sequence length distribution after merging reads, for one library. The plot is meant to help making a sensible choice for sequence length filtering. **(B)** Number of sequences per sample. This histogram is part of the OTU report (including all libraries). **(C)** Number of sequences after individual preprocessing steps. ‘Assembled’ refers to the number of read pairs which could be merged based on their overlap. This plot is part of the library report. **(D)** Number of sequences after individual steps after potentially combining several libraries (total number of reads) and generating OTUs. ‘Derep.’ indicates the number of dereplicated reads. ‘OTUs’ is the total number of OTUs and ‘Assigned OTUs’ the number of OTUs with a taxonomic assignment. ‘No singletons’ refers to the number of OTUs excluding singleton OTUs and ‘Assigned NO singletons’ the number of singleton-free OTUs with a taxonomic assignation. The plot is part of the OTU report. **(E)** Krona chart for one sample. The krona charts are interactive and can be viewed with a web browser. Colors indicate the taxonomic groups that the OTU was assigned to. Each ring of the pie chart represents a different taxonomic level. An example of a full library report is shown in supplemental file 2, and an OTU report is provided in supplemental file 3.

Next, *Cascabel* will cluster reads from the sequence data of all libraries into OTUs. We used 97% sequence identity with uclust to generate roughly 2.5 million OTUs. We chose the longest sequence of an OTU as representative sequence to be used for taxonomic placement of the OTU. Alternative OTU and representative sequence picking methods provided by *Cascabel* are listed in Table 1. Then we used VSEARCH to assign taxonomy to the representative sequences based on the SILVA database (SILVA version 132).

From the abundances of the OTU sequences within each of the samples, *Cascabel* creates an OTU abundance table. The OTUs can further be grouped at higher taxonomic levels depending on the desired resolution. Subsequently, the user can opt to remove singletons, align representative sequences, filter the alignment and make a phylogenetic tree. Removing singletons reduced the number of OTUs in the analyzed dataset to roughly 500.000. To align representative sequences, *Cascabel* offers pynast (25), mafft (26), infernal (27), clustalw (28), and muscle (29), and we used pynast on our data. A phylogenetic tree can be generated with muscle, clustalw, raxml (30) and fasttree (31), and we applied fasttree (Table 1).

Finally, *Cascabel* generates HTML and optionally pdf reports of the analyses, documenting all software and parameters used. If more than one library was analyzed, there will be a report for each library as well as a report summarizing all libraries (otu_report). Among other graphics, the otu_report shows the percentages and the total number of reads after filtering (‘combined reads’), dereplicated reads, OTUs, OTUs assigned to a taxonomic level, OTUs excluding singletons (‘no singletons’), and assigned OTUs excluding singletons. The graph for the analyzed example data is shown in Figure 2D. Supplementary files 2 and 3 show the library and the otu_report for the demonstration data, respectively. In addition, *Cascabel* generates an interactive Krona chart (32) for the run which displays community composition for individual samples or the complete data set. The Krona chart shows the taxonomic assignments in an interactive HTML document composed of a multi-layered pie-chart and the user can zoom and browse these different levels. An example is shown in Figure 2E.

The user can make use of all intermediate files generated by individual rules, and most importantly the OTU table and representative sequences for follow-up analyses. To save disk space, the user can also opt to have *Cascabel* remove temporary files at the end of the analyses. For many rules, the user can pass additional parameters to the command or tool at hand using the ‘extra_params’ parameter in the config file.

## Discussion

*Cascabel* has been developed at the Royal Netherlands Institute for Sea Research (NIOZ) to facilitate, unify and easily track data provenance of amplicon sequence data analyses. Apparent advantages of using this pipeline compared to custom scripts are that the individual steps of the pipeline have been tested by many members of the community at the NIOZ who are experienced in amplicon sequencing data analyses (34–37), and therefore should contain fewer mistakes than scripts that were written for a specific analysis by one person. Moreover, community knowledge and experience have created a workflow which is probably more comprehensive and powerful than one that was created by a single person. In addition, the availability of the pipeline has facilitated comparing and integrating research results from different data sets generated at the NIOZ because scientists can agree on certain settings and reference database versions and the pipeline guarantees that the analyses are performed in the same way. Because *Cascabel* keeps track of data provenance, documenting the process of analyzing the data to generate results, it also facilitates preparing research manuscripts. While most of the scientific journals request the raw sequencing data to be submitted to a public repository for many years already, also reporting data provenance becomes more important. The journal ‘Nature’, for example, requires authors to make materials, data, code, and associated protocols available (38). *Cascabel* facilitates providing data, code and protocols. Public sequence repositories often require the raw data to be submitted per sample, but sample demultiplexing typically takes place after merging read pairs such that the raw data cannot be recovered. Therefore *Cascabel* demultiplexes the raw data in parallel to the analyses such that it is ready for public data repository deposition. The code of Cascabel is open source and all analyses are protocoled in the reports and config file, complying with the rules for reproducible computational research described by Sandve et al. (39).

DNA sequencing technology, algorithms and analysis approaches are constantly evolving. It is logical that pipelines lag behind with the most recent developments because it takes time to test and integrate new modules. Because *Cascabel* is a Snakemake workflow, it is flexible and easy to extend to encompass more or alternative rules. We are constantly working on extending the range of applications and making alternative approaches, like generating amplicon sequence variants (ASVs) instead of OTUs, available. *Cascabel* provides reference databases for taxonomy assignment and chimera detection, but the user can always supply a different database and specify that in the config file. Moreover, *Cascabel* is not limited to Illumina sequence data that we used for demonstration purposes, but can handle sequence data from other technologies which produce short reads from amplicons as well (e.g. Ion Torrent). With some minor modifications, *Cascabel* can even be used to analyze long read amplicon sequence data.

Galaxy (40) might be a user-friendly web-based alternative to *Cascabel* which offers interfaces to VSEARCH and mothur executables. Having a medium-sized user group at the institute, we did not want to overload a public server and setting up and maintaining our own server would also need resources that we preferred to allocate to the development of a workflow for which we have full control and flexibility. With *Cascabel* being invoked from the command line, the user can make use of the full potential that Snakemake has to offer, e.g., `-prioritize` to force the execution of specific rules prior to others when distributing tasks across computing resources, `-until` to run the pipeline up to a specific rule, `-summary`, which shows the rules executed so far and `-dag` which shows the rules executed and the ones yet to be done in a directed acyclic graph. Moreover, we consider *Cascabel*’s report an essential element to move forward in terms of user-friendly data provenance and reproducibility.

We have presented *Cascabel*, an open source pipeline to analyze amplicon sequence data based on the workflow engine Snakemake. The pipeline can be easily installed via conda, comes with documentation and a wiki on github and can be executed by users with basic command line skills. At the same time, *Cascabel* is flexible, offering alternative options for most of the steps and supporting custom reference databases, and can easily be modified and extended by users with computational skills. We believe that *Cascabel* will prove to be useful to scientist who need more flexibility and throughput than provided by tools based on web servers, but do not want to or cannot generate their own command-line based workflow.

## Methods

### Sampling and DNA extraction

Suspended particulate matter (SPM) was collected from Lake Chala from September 2013 to May 2014 from a total of 111 samples as described by (41). DNA was extracted from 1/32 section of the filters on which SPM was collected by using the PowerSoil DNA extraction kit (Mo Bio Laboratories, Carlsbad, CA, USA).

### DNA sequencing

The V4 region of the 16S rRNA gene were amplified with the primers forward:

515F (Parada): GTGYCAGCMGCCGCGGTAA (42) and reverse:

806R (Apprill): GGACTACNVGGGTWTCTAAT (43). We made use of 12 nucleotide Golay barcodes at the beginning of the forward read. Paired-end sequencing of 250 nt was performed on an Illumina MiSeq instrument (Illumina, San Diego, CA) using the Truseq DNA nano LT kit for library preparation and V3 sequencing chemistry at the sequencing facility of the University of Utrecht (USEQ), the Netherlands. The data is publically available at NCBI, BioProject PRJNA526242.

### Sequence analysis

All the settings and parameters chosen to analyze the example data set are given in the config file (supplementary file 1) and the reports (supplementary files 2 and 3).

## Author Contributions

AA implemented the pipeline, with contributions from JCE. MB, LV, SB and HW tested the pipeline. AA, JvB and JCE designed the pipeline, with contributions from LV. JCE wrote the manuscript. All authors contributed to and approved the final version of the manuscript.

## Acknowledgments

We thank Hans Malschaert for IT support and the bioinformatics community at the NIOZ for comments and suggestions about the pipeline. We acknowledge S. Vreugdenhil and M. Brouwer for technical and analytical support. Fieldwork with collection of the studied sample materials was carried out with permission from the government of Kenya through permit 13/001/11C to D.V. In accordance with National Environmental Management Authority regulations in the context of the Nagoya Protocol, DNA extracts of the analyzed suspended-particulate samples are archived at the National Museums of Kenya (NMK), under voucher numbers NMK:BCT:80001 to NMK:BCT:80221.

