## Supplemental File 1 for "*Cascabel*: a flexible, scalable and easy-to-use amplicon sequence data analysis pipeline": SuppFile1-config_NIOZ66.yaml

```
################################################################################
#                             CONFIGURATION FILE                               #
#------------------------------------------------------------------------------#
# Configuration file for  CASCABEL pipeline.                                   #
# Set the parameters below, save the file and run Snakemake.                   #
# The file format is yaml (http://www.yaml.org/) In this file, you specify your#
# input data, barcode mapping file and you can choose tools and parameters     #
# according to your needs. Most rules and parameters have default settings.    #
# It is very important to keep the indentation of the file (don’t change the   #
# tabs and spaces), as well as the name of the parameters/variables. But you   #
# can of course change the values of the parameters to deviate from the default# 
# settings. Any text after a hashtag (#) is considered a comment and will be   #
# ignored by Snakemake.                                                        #
#                                                                              #
# @Author: Julia Engelmann and Alejandro Abdala                                #
# @Last update: 31/07/2019                                                     #
################################################################################

################################################################################
#                       GENERAL PARAMETERS SECTION                             # 
#------------------------------------------------------------------------------#
# The general parameters section defines parameters that are global or general #
# for the complete workflow.                                                   #
################################################################################

#------------------------------------------------------------------------------#
#                             Execution mode                                   #
#------------------------------------------------------------------------------#
# This parameter allows the user to inspect intermediate files in order to     #
# finetune some downstream analises, re-do previous steps or exit the workflow.#	            
# -interactive   Set this flag to "T" (default) in order to interact at some   #
#                specific steps with the pipeline. "F" will try to run all the #
#                pipeline without communicating intermediate results until the #
#                report.                                                       # 
# For a list for all the interactive checkpoints take a look at the following  #
# link: TBD                                                                    #
#------------------------------------------------------------------------------#
interactive : "F"

#------------------------------------------------------------------------------#
#                             Project Name                                     #
#------------------------------------------------------------------------------#
# The name of the project for which the pipeline will be executed. This should #
# be the same name used as the first parameter with the init_sample.sh script  #
# (if used for multiple libraries).                                            #
#------------------------------------------------------------------------------#
PROJECT: "Report_NIOZ66"

#------------------------------------------------------------------------------#
#                            LIBRARIES/SAMPLES                                 #
#------------------------------------------------------------------------------#
# SAMPLES/LIBRARIES you want to include in the analysis.                       #
# Use the same library names as with the init_sample.sh script.                #
# Include each library name surrounded by quotes, and comma separated.         #
# i.e LIBRARY: ["SAMP_1","SAMP_2",..."SAMP_N"]                                 #
#------------------------------------------------------------------------------#
LIBRARY: ["NIOZ66"]

#------------------------------------------------------------------------------#
#                               RUN                                            #
#------------------------------------------------------------------------------#
# Name of the RUN - Only use alphanumeric characters and don't use spaces.     #
# This parameter helps the user to execute different runs (pipeline executions)#
# with the same input data but with different parameters (ideally).            #
# The RUN parameter can be set here or remain empty, in the latter case, the   #
# user must assign this value via the command line.                            #
# i.e:  --config RUN=run_name                                                  #
#------------------------------------------------------------------------------#
RUN: "final_report_example"

#------------------------------------------------------------------------------#
#                           Description                                        #
#------------------------------------------------------------------------------#
# Brief description of the run. Any description written here will be included  #
# in the final report. This field is not mandatory so it can remain empty.     #
#------------------------------------------------------------------------------#
description: "This is the report generated by CASCABEL"

#------------------------------------------------------------------------------#
#                             INPUT TYPE                                       #
#------------------------------------------------------------------------------#
# CASCABEL supports two types of input files, fastq and gzipped fastq files.   #     
# This parameter can take the values "T" if the input files are gzipped         #
# (only the reads!, the metadata file always needs to be uncompressed) or "F"  #
# if the input files are regular fastq files.                                   #
#------------------------------------------------------------------------------#
gzip_input: "F"

#------------------------------------------------------------------------------#
#                             INPUT FILES                                      #
#------------------------------------------------------------------------------#
# There are two ways to initialize the required project structure in order to  #
# run the pipeline. The first one is to use the script init_sample.sh before   #
# running the pipeline; more info at:                                          #
# https://github.com/AlejandroAb/CASCABEL/wiki#initialize-structure-for-multiple-libraries 
# If you are planning to run the pipeline for one single library, you can      #
# supply all the necessary input files directly into this section and avoid    #
# running the previous mentioned script.                                       #
# Here you have to enter the FULL PATH for both: the raw reads and the metadata# 
# file (barcode mapping file). The metadata file is only needed if you want to #
# perform demultiplexing.                                                      #
# - fw_reads:  Full path to the raw reads in forward direction (R1)            #
# - rw_reads:  Full path to the raw reads in reverse direction (R2)            #
# - metadata:  Full path to the metadata file with barcodes for each sample    #
#              to perform library demultiplexing                               #
#------------------------------------------------------------------------------#
fw_reads: "/export/data/villanueva/Documents_Laura/NIOZ66/NG-10020_NIOZ_SG1_lib132214_4719_1_1.fastq"
rv_reads: "/export/data/villanueva/Documents_Laura/NIOZ66/NG-10020_NIOZ_SG1_lib132214_4719_1_2.fastq"
metadata: "/export/data/villanueva/Documents_Laura/NIOZ66/nioz66_map.txt"


################################################################################
#                          REPORT PARAMETER SECTION                            #
#------------------------------------------------------------------------------#
# This section defines parameters that will influence the type of report to be #
# generated at the end of the workflow.                                        #
################################################################################


#------------------------------------------------------------------------------#
#                           PDF Report                                         #
#------------------------------------------------------------------------------#
# By default, CASCABEL creates the final report in HTML format. In order to    #
# create the report also as pdf file, set this flag to "T".                    #
# Important! in order to convert the file to pdf format, it is necessary to    #
# execute the pipeline within a Xserver session i.e MobaXterm or ssh -X.       #
# One way to validate if your active session is using an Xserver, execute      #
# 'echo $DISPLAY'  on a command line terminal. If this returns empty, you do   #
# not have an Xserver session.                                                 #
# - pdfReport            "T" for generate pdf report or "F" to skip it.        #
# - wkhtmltopdf_command  name w/wo path to execute the html to pdf translation #
#------------------------------------------------------------------------------#
pdfReport: "T"
wkhtmltopdf_command: "wkhtmltopdf"

#------------------------------------------------------------------------------#
#                           Portable Report                                    #
#------------------------------------------------------------------------------#
# CASCABEL creates the final report in HTML format, containing references to   #
# other images or links. Therefore just copying the HTML files for sharing or  #
# inspecting the results will break these links.                               #
# By setting 'portableReport' to true "T", CASCABEL will generate a zip file   #
# with all the resources necessary to share and distribute CASCABEL's report.  #
#------------------------------------------------------------------------------#
portableReport: "T"

#------------------------------------------------------------------------------#
#                           Krona Report                                       #
#------------------------------------------------------------------------------#
# Krona allows hierarchical data to be explored with zooming, multi-layered pie#
# charts. The interactive charts are self-contained and can be viewed with any #
# modern web browser.                                                          #
# - report        Indicate with "T"/"F" if CASCABEL should generate a Krona    #
#                 chart.                                                       #
# - ktImportText  Name of the command to invoque this krona utility.           #
# - samples       Indicate the samples to be included in the chart, use comma  #
#                 separated values of samples (same name as the ones supplied  #
#                 at the metadata barcode file). Or "all" to include all the   #
#                 samples.                                                     #
# - otu_table     Target OTU table for the report. Use "default" to use the    #
#                 filtered OTU table (exclude singletons). Or "singletons" to  #
#                 use the non filtered OTU table (include singletons)          #  
# - extra_params  Any other extra parameter from ktImportText tool. default    #
#                 "-n root_extra"                                              #
#------------------------------------------------------------------------------#

krona:
  report: "T"
  command: "ktImportText"
  samples: "all"
  otu_table: "default"
  extra_params: "-n root_extra"


################################################################################
#                        Specific Parameters Section                           #
#------------------------------------------------------------------------------#
# In this section of the configuration file, you can find all the parameters   #
# used to run the different rules during the execution of the pipeline.        #
# Some of the entries below contain a parameter called "extra_params".         #
# This parameter is designed to allow the user to pass any other extra         #
# parameter to the program invoked by the rule, as some rules do not list all  #
# the parameters of the underlying tool explicitly. In these cases, the user   #
# can specify any other parameter using "extra_params".                        #  
################################################################################

#------------------------------------------------------------------------------#
#                        Quality control with FastQC                           #
# rules: fast_qc, validateQC                                                   #
#------------------------------------------------------------------------------#
# FastQC evaluates 12 main concepts on the sequences: basic statistics, per    #
# base sequence quality, per tile sequence quality, per sequence quality       #
# scores, per base sequence content, per sequence GC content, per base N       #
# content, sequence length distribution, sequence duplication levels,          #
# overrepresented sequences, adapter content and Kmer content.                 #
# - command       Command needed to invoke fastqc [default: "fastqc"].         #
# - extra_params  Extra parameters. See fastqc --help for more info.           #
# - qcLimit       Set the maximum number of FastQC FAILS (from the 12 tests    #
#                 evaluated) accepted before interrupting the workflow if the  #
#                 'interactive' mode is equal to "T"                           #
#------------------------------------------------------------------------------#
fastQC:
  command: "fastqc"
  extra_params: ""
  qcLimit : 3

#------------------------------------------------------------------------------#
#           Assemble fragments (merge forward with reverse reads)              #
# rule: pear                                                                   #
#------------------------------------------------------------------------------#
# This step is performed to merge paired reads.                                #
# - t             Minimum length of reads after trimming the low.              #
# - v             Minimum overlap size. The minimum overlap may be set to 1    #
#                 when the statistical test is used. (default in pear: 10)     #
# - j             Number of threads to use.                                    #
# - p             The p-value cutoff used in the statistical test. Valid       #
#                 options are: 0.0001, 0.001, 0.01, 0.05 and 1.0. Setting 1.0  #
#                 disables the test. (default: 0.01)                           #
# - extra_params  Extra parameters. See pear --help for more info.             #
# - prcpear       The minimun percentage of expected peared reads, if the      #
#                 actual percentage is lower and the 'interactive' parameter   #
#                 is set to "T";  a warning message will be shown.             #
#------------------------------------------------------------------------------#
pear:
  command: "pear"
  t: 100
  v: 10
  j: 6
  p: 0.05
  extra_params: ""
  prcpear: 90

#------------------------------------------------------------------------------#
#                  FastQC on merged/assembled fragments                        #
# rule: fastQCPear                                                             #
#------------------------------------------------------------------------------#
# Once the paired end reads have been merged into one fragment, run FastQC     #
# again to check their quality. Set this option to "T" (true) or "F" (false)   #
# in order to execute or skip this step.                                       #
#------------------------------------------------------------------------------#
fastQCPear: T

#------------------------------------------------------------------------------#
#                                  QIIME                                       #
# rule: bc_mapping_validation, extract_barcodes, extract_barcodes_unassigned,  #
# split_libraries, split_libraries_rc, search_chimera, cluster_OTUs,           #
# pick_representatives, assign_taxonomy, make_otu_table, summarize_taxa,       #
# filter_rep_seqs, align_rep_seqs, filter_alignment, make_tree                 #
#------------------------------------------------------------------------------#
# Different QIIME scripts are used along the pipeline and in order to execute  #
# these scripts, they need to be located on the user PATH environmental        #
# variable, or QIIME's bin directory need to be included in the parameter:     #
# 'path' below. This parameter will be used by the pipeline for all the rules  #
# that use a Qiime script.                                                     #
#------------------------------------------------------------------------------#
qiime:
  path: ""

#------------------------------------------------------------------------------#
#                                    R                                         #
# rules:  correct_barcodes, correct_barcodes_unassigned, histogram_chart       #
#------------------------------------------------------------------------------#
# R is used by different rules whitin the pipeline. In order to run these      #
# rules, CASCABEL uses the Rscript command.                                    #
# Here you can change the command to call Rscript.                             #
# - path     Rscript command, [default "Rscript"].                             #
#------------------------------------------------------------------------------#
Rscript:
  command: "Rscript"

#------------------------------------------------------------------------------#
#                                  JAVA                                        #
# rules:  write_dmx_files, degap_alignment, remap_clusters                     #
#------------------------------------------------------------------------------#
# Java is used by different rules whitin the pipeline. In order to run java,   #
# the pipeline needs to know how to invoke it.                                 #
# Here you can change the command to invoke java.                              #
# - path     java command, [default: "java"].                                  #
#------------------------------------------------------------------------------#
java:
  command: "java"


#------------------------------------------------------------------------------#
#                   Demultiplex input files                                    #
# rule: write_dmx_files                                                        #
#------------------------------------------------------------------------------#
# CASCABEL optionally performs library demultiplexing for barcoded reads.      #
# This feature can be turned ON/OFF with the following options:                #
# - demultiplex         "T". If the pipeline is going to demultiplex the input #
#                       files. In this case, the metadata file has to be       #
#                       provided.                                              #
#                       "F". If the input files are already demultiplexed.     #
# - create_fastq_files  "T" or "F". If 'demultiplex' = T and this is also T,   #
#                       the pipeline will create demultiplexed fastq files per #
#                       sample.                                                #
# - dmx_params:         Parameters to pass on to the demultiplexing script.    #
#                       For example, the user can change the prefix and suffix #
#                       of the output files. To see the available parameters,  #
#                       run: java -cp Scripts DemultiplexQiime                 #
#------------------------------------------------------------------------------#

demultiplexing:
  demultiplex: "T"
  create_fastq_files: "T"
  dmx_params: ""

#------------------------------------------------------------------------------#
#                           Extract barcodes                                   #
# rules: extract_barcodes, extract_barcodes_unassigned                         #
#------------------------------------------------------------------------------#
# These rules extracts barcodes from the reads, and at the end generates two   #
# files: one with all the barcodes and a second one with the sequences without #
# the barcodes.                                                                #
# - c             This parameter allows to choose the barcode configuration:   #
#                 "barcode_single_end". The reads starts with the barcode      #
#                 sequence.                                                    #
#                 "barcode_paired_stitched". Input has barcodes at the         #
#                 beginning and end of the read.                               #
#                 "barcode_paired_end". This option is not valid here since    #
#                 the reads have already been merged to one fragment.          #
# - bc_length     If 'c' is "barcode_paired_stitched" use both:                #
#                 --bc1_len X and --bc2_len Y.                                 #
#                 If 'c' is "barcode_single_end" use only --bc1_len X.         #
#                 Where X and Y are the lengths of the barcodes.               #
# - extra_params  Extra parameters. See extract_barcodes.py -help.             #
#------------------------------------------------------------------------------#

ext_bc:
  c: "barcode_single_end"
  bc_length: "--bc1_len 12"
  extra_params: ""

#------------------------------------------------------------------------------#
#                             Barcode correction                               #
# rules: correct_barcodes, correct_barcodes_unassigned                         #
#------------------------------------------------------------------------------#
# This parameter allows error correction of barcodes. First, barcodes perfectly#
# matching a sample barcode in the mapping file will be assigned to the        #
# samples. If error correction is enabled, barcodes will be assigned to the    #
# closest barcode with increasing number of mismatches until the maximum number# 
# of mismatches has been reached, e.g. if the mismatch is equal to "2", first  #
# barcodes with one mismatch will be assigned, then barcodes with 2 mismatches.#
#    bc_mismatch:  Number of allowed missmatches.                              # 
#                  bc_mismatch = 0: don't allow mismatches in the barcode.     #
#                  bc_mismatch > 0: correct bc_mismatch bases at maximum.      #
#------------------------------------------------------------------------------#
bc_mismatch: 2

#------------------------------------------------------------------------------#
#                             Split libraries                                  #
# rule: split_libraries, split_libraries_rc                                    #
#------------------------------------------------------------------------------#
# These rules performs demultiplexing of Fastq sequence data where barcodes and#
# sequences are contained in two separate fastq files.                         #
# - q             Maximum unacceptable Phred quality score. e.g., for Q20 and  #
#                 better, specify -q 19). [default: 19]                        #
# - r             Maximum number of consecutive low quality base calls allowed #
#                 before truncating a read. [default: 3]                       #
# - barcode_type  The type of barcode used. This can be an integer, e.g., "6"  #
#                 or “golay_12” for golay error-correcting barcodes.           #
# - extra_params  Any extra parameter. Run split_libraries_fastq -h to see all #
#                 options.                                                     #
#------------------------------------------------------------------------------#
split:
  q: "19"
  r: "5"
  barcode_type:  "12"
  extra_params: ""

#------------------------------------------------------------------------------#
#                         Align reads vs a reference database                  #
# rule: align_vs_reference                                                     #
#------------------------------------------------------------------------------#
# In some cases you may want to align the reads against a reference database   #
# before generating OTUs. This facilitates removing technical sequence	       #
# (primers, adapters) at the beginning and/or end of the reads, filters	       #
# potential chimeric sequences or sequences of no interest, which the primers  #
# amplified but which are not part of the study (e.g. reads of human origin).  #
# Therefore, this alignment step can improve the OTU clustering and further    #
# taxonomy assignation. In order to run the alignment here, use 'align': "T"   #
# and bear in mind that whenever you do this, you should only do it with small #
# to medium sized data bases, because sequence alignment is computationally    #
# costly.                                       	    		       #
# - mothur_cmd:      Enter the command to call mothur [default: "mothur"].     #
# - align:           "T" or "F" to run or skip this rule respectively.         #
# - dbAligned:       Database to perform the alignment with.                   #
# - cpus:            Number of CPUs to perform the alignment with.             #
#------------------------------------------------------------------------------#
align_vs_reference:
  mothur_cmd: "mothur"
  align: "F"
  dbAligned: ""
  cpus: 4

#------------------------------------------------------------------------------#
#                             Remove adapters                                  #
# rule: cutadapt                                                               #
#------------------------------------------------------------------------------#
# This rule runs Cutadapt. Cutadapt searches for adapters in the reads and     #
# removes them when it finds any. Unless you use a filtering option, all       #
# reads that were present in the input file will also be present in the output #
# file, some of them trimmed, some of them not.                                #
# For details, see: http://cutadapt.readthedocs.io/en/stable/guide.html        #
# - cutAdapters  if T (true) remove adapters, if F do not remove adapters      #
#                [default: F].                                                 #
# - command      Necessary command to invoque cutadapt [default: "cutadapt"].  #
# - adapters     "-a ADAPTER" Removes adapters at the the 3' end of the        #
#                sequence. Cutadapt deals with 3' adapters by removing the     #
#                adapter and any sequence that may follow. Add the "$"         # 
#                character to the end of an adapter sequence in order to anchor#
#                the adapter to the end of the read, such as "-a ADAPTER$". The#
#                adapter will only be removed if it is a suffix of the read.   #
#                "-g ADAPTER" Removes adapters at the 5' end. If you want to   #
#                trim only if the sequence starts with the adapter, use        #
#                "-g ^ADAPTER". The "^" character indicates that the adapter is#
#                'anchored' at the beginning of the read. In other words: The  #
#                adapter is expected to be a prefix of the read.               #
#                If your sequence of interest is 'framed' by a 5' and a 3'     #
#                adapter and you want to remove both adapters, then you may    #
#                want to use a linked adapter. A linked adapter combines an    #
#                anchored 5' adapter and a 3' adapter. The 3' adapter can be   #
#                regular or anchored. The idea is that a read is only trimmed  #
#                if the anchored adapters occur. Thus, the 5' adapter is always# 
#                required, and if the 3' adapter was specified as anchored, it #
#                also must exist for a successful match, e.g.                  #
#                -a GTGYCAGCMGCCGCGGTAA...ATTAGAWACCCVNGTAGTCC                 #
# - extra_params Any extra parameter. A useful extra parameter here is         #
#                --match-read-wildcards which interprets IUPAC wildcards in    #
#                reads.                                                        # 
# More info and options at http://cutadapt.readthedocs.io/en/stable/guide.html #
#------------------------------------------------------------------------------#
cutAdapters: "F"
cutadapt:
  command: "cutadapt"
  adapters: ""
  extra_params: "--match-read-wildcards"

#------------------------------------------------------------------------------#
#                         Identify chimeric sequences                          #
# rule: search_chimera                                                         #
#------------------------------------------------------------------------------#
# This rule will be executed if and only if the option 'search' is set to "T"! #
# Chimeric sequencesa are predicted using usearch61. This algorithm performs   #
# both de novo (abundance based) chimera and reference based detection.        #
# Unclustered sequences should be used as input rather than a representative   # 
# sequence set, as the sequences will be clustered by usearch61 to get         #
# abundance data. The results are all input sequences not flagged as chimeras. # 
# For details, see: http://drive5.com/usearch/usearch_docs.html                #
# It is possible to use two other methods: the Blast_fragments approach and    #
# ChimeraSlayer, however we recomend to use usearch61 since it uses unclustered#
# sequences unlike the other two algorithms, which use representative          #
# sequences.                                                                   # 
# If you are using usearch61, bear in mind that you can use a reference        #
# database via extra_params, e.g., "-r /path/to/gold_db/gold.fa"               #
# - method        This method could be any of the ones used by the script      #      
#                 identify_chimeric_seqs, but because "usearch61" is the most  #
#                 time-efficient, for the moment CASCABEL only provides        #
#                 "usearch61".                                                 #
# - threads       Number of threads to use.                                    #
# - extra_params  Any extra parameter. It is recomended to run the chimeric    #
#                 search against a chimera database, e.g.,                     #
#                 "-r /export/data/databases/gold_db/gold.fa"                  #
# - search        "T" | "F" (true or false) to execute chimera checking or not.#
#------------------------------------------------------------------------------#
chimera:
  search: "F"
  method: "usearch61"
  threads: 6
  extra_params: ""

#------------------------------------------------------------------------------#
#                          Length filtering                                    #
# rule: remove_short_long_reads                                                #
#------------------------------------------------------------------------------#
# This rule executes a python script in order to filter the reads based on     #
# their length.                                                                #
# First, this script generates a histogram based on the read lengths           #
# distribution. Next, this histogram is used by the same script in different   # 
# ways depending on the pipeline's execution mode (interactive or automatic).  #
# If the pipeline is executed in "Interactive" mode, it will always stop at    #
# this step and let the user choose between the following options:             #
#      * Use the values specified in the configuration file (the ones from the #
#        parameters specified here, 'longs' and 'shorts').                     #
#      * Filter the reads based on the median of the sequence length           #
#        distribution +/- an offset value.                                     #
#      * Do not filter any sequence.                                           #
#      * Stop the pipeline.                                                    #
# If the pipeline is executed in non-interactive/automatic mode, the pipeline  #
# will not stop and the reads will be filtered according to the                #
# 'non_interactive_behaviour' value.                                           #
# - non_interactive_behaviour  Behaviour for the non-interactive/automatic     # 
#                              mode. Valid options are:                        #  
#                              * CFG: use the values from the configuration    #
#                                     file ('longs' and 'shorts').             #
#                              * AVG: use the values from the median           #
#                                     distribution.                            #           
#                              * NONE: do not filter any read.                 #
#                              [default: CFG]                                  #
# - offset                     value used to determine the bounds for the      #
#                              filtering when 'interactive' is set to 'F' and  #
#                              non_interactive_behaviour is equal to 'AVG'.    # 
#                              [default: 10].                                  #
# - longs                      Maximum read length.                            #
# - shorts                     Minimumin read length                           #
#------------------------------------------------------------------------------#
# REMARK: If your library contains more than one expected fragment length, you #
# can either:                                                                  #
# A) Do not filter any length.                                                 #
# B) Use inclusive boundaries.                                                 #
# C) Rerun the pipeline for all the expected fragment lengths, e.g.,           #
#    "--forcerun remove_short_long_reads". In this case, do not forget to      #
#    backup previous results, otherwise, they will be overwritten.             #
#------------------------------------------------------------------------------#
rm_reads:
  non_interactive_behaviour: "AVG"
  offset: 15
  longs: 1000
  shorts: 10


#------------------------------------------------------------------------------#
#                               Dereplicate                                    #
# rule: dereplicate, pick_derep_representatives                                #
#------------------------------------------------------------------------------#
# This parameters allows the user to dereplicate the sequences over the FULL   #
# LENGTH (100% identity) before applying any OTU picking starategy. This is    #
# adviced for very large datasets, when OTU picking methods take too long or   #
# have memory issues.                                                          #
# - dereplicate    Dereplicate sequences over their full length (F/T)          #
#                  [default: "F"].                                             #
# - vsearch_cmd    Command for calling vsearch [default: "vsearch"].           #
# - min_abundance  Minimum abundance for output from dereplication.            #
# - strand:        plus|both, search "plus" or "both" strands                  #
#                  [default: "both"].                                          #
# - extra_params   Dereplication is performed using vsearch, you can add       #
#                  different options described by vsearch --help.              #
#------------------------------------------------------------------------------#
derep:
  dereplicate: "T"
  vsearch_cmd: "vsearch"
  min_abundance: 1
  strand: "both"
  extra_params: ""

#------------------------------------------------------------------------------#
#                               OTU picking                                    #
# rule: cluster_OTUs                                                           #
#------------------------------------------------------------------------------#
# The OTU picking step assigns similar sequences to operational taxonomic      #
# units, or OTUs, by clustering sequences based on a user-defined similarity   #
# threshold. Sequences which are similar at or above the threshold level are   #
# taken to represent the presence of a taxonomic unit (e.g., approximately at  #
# genus level, when the similarity threshold is set at 0.94) in the sequence   #
# collection. Swarm takes a different clustering approach which does not       #
# require setting a threshold. Instead, clusters are formed using sequence     # 
# graphs and abundance.                                                        #
# - s              Sequence similarity threshold between 0 and 1. This applies #
#                  for the following methods 'm': uclust, uclust_ref, usearch, #
#                  usearch_ref, usearch61,  usearch61_ref, sumaclust, and      #
#                  sortmerna. [default:0.97].                                  #
# - m              OTU picking method. Valid choices are: sortmerna, mothur,   #
#                  trie, uclust, uclust_ref, usearch, swarm.                   #
# - extra_params   Any extra parameter. Run 'pick_otus.py -h' to see all       #
#                  options.                                                    # 
#------------------------------------------------------------------------------#
pickOTU:
  s: "0.97"
  m: "uclust"
  cpus: "6"
  extra_params: ""

#------------------------------------------------------------------------------#
#                        Select representative sequences                       #
# rule: pick_representatives                                                   #
#------------------------------------------------------------------------------#
# After picking OTUs, this rule picks a representative sequence for each OTU.  #
# - m             Method for picking representative sequences. Valid choices   #
#                 are: random, longest, most_abundant, first [default: "first"]# 
#                 Note: "first" chooses the cluster seed when picking otus with#
#                 uclust.                                                      #
# - extra_params  Any extra parameter. Run 'pick_rep_set.py -h' to see all     #
#                 options.                                                     #
#------------------------------------------------------------------------------#
pickRep:
  m: "longest"
  extra_params: ""

################################################################################
#                            Assign taxonomy                                   #
# rule: assign_taxonomy                                                        #
#------------------------------------------------------------------------------#
# Performs taxonomy assignment for the representative sequences.               #
# This step can be performed using three different tools (parameter 'tool'):   #
# 1) VSEARCH.  Compare target sequences 'db_file' to the query sequences to    #
#              assign taxonomy, using global pairwise alignment.               #
# 2) BLAST.    This uses BLAST+.                                               #
# 3) QIIME.    In this case, CASCABEL runs the assign_taxonomy.py script which #
#              can use any of the following methods:                           #
#  3.1) BLAST.  To use blast via QIIME, use "blast" as 'method' below.         # 
#               This uses the old version of BLAST (not BLAST+).               #
#               This method can work with either a BLAST database, setting     #
#               'dbType' to "-b" and 'dbFile' with the full path to a BLAST    #
#               database, or with a fasta file with 'dbTYpe' set to "-r" and   #
#               'dbFile' pointing to the fasta file (full path must be used).  #
#  3.2) UCLUST. A method based on sequence clustering. To use this method type #
#               "uclust" as 'method' below. This method ONLY works with        #
#               'dbType' set to "-r" and 'dbFile' full path to a fasta file.   #
#  3.3) RDP.    The Ribosomal Database Project (RDP) Classifier, a naive       #
#               Bayesian classifier, can rapidly and accurately classify       #
#               bacterial 16S rRNA sequences. To use this method, use "rdp" as #
#               'method' below.                                                #
#               NOTE: In order to run RDP it is necessary to provide the       #
#               classifier path. To do so, please include the following line   #
#               on the "extra_params":                                         #
#               '--rdp_classifier_fp /path/to/rdp_classifier-2.2.jar'          #
#               Due to some compatibility issues, RDP needs a custom fasta file#
#               and taxonomy mapping file. You can find the appropriate files  # 
#               at the following references:                                   #
#               'dbFile' "/gg/gg_13_8_otus/rep_set/97_otus.fasta"              #
#               'mappFile' "/gg/gg_13_8_otus/taxonomy/97_otu_taxonomy.txt"     #
#            These 3 different methods can use different sets of options in    #
#            order to fine tune the taxonomy assignment. These options can be  #
#            realized with the extra_params parameter. To see what options are #
#            available for each 'method', type on the command line             #
#            'parallel_assign_taxonomy_<method>.py -h'.                        #
# - tool     Which tool to use, options are "vsearch", "qiime" and "blast".    #
# ----blast parameters                                                         #
#    - command         Command to invokw blastn.  [default: "blastn"].         #
#    - blast_db        Full path to target BLAST databse for the assignation.  #
#    - fasta_db        Full path to a fasta file with sequences for the        #
#                      assignation. In case of providing both (a Blast database#
#                      and a fasta file, Cascabel will give priority to the    #
#                      database 'blast_db'.                                    #
#    - mapFile         Mapping file between sequence accessions and their      #
#                      taxonomy. This file should be a two column file, tab    #
#                      separated with the accessions in the first column and   #
#                      the taxonomy in the second column with the taxonomic    #
#                      levels separated by 'taxo_separator', e.g.              #
#                      'AY190.44  Bacteria;Firmicutes;Bacilli;...'             # 
#    - taxo_separator  Character used to split taxonomic levels specified at   #
#                      the taxonomy mapping file 'mapFile'. [default: "';'"]   #
#    - max_target_seqs Maximum number of target sequences per sequence query.  #
#                      All the target sequences are taking into account and    #
#                      their taxonomies are mapped to their Lowest Common      #
#                      Ancestor (LCA) with the stampa approach	   	       #
#		       (https://github.com/frederic-mahe/stampa).	       #
#		       "1" disables LCA mapping.                               #
#    - jobs            Number of cpus.                                         #
#    - identity        Minimum percentage of identity (between 0 and 1).       #
#    - extra_params    Any extra parameter. Run 'blastn -help' to see all      #
#                      options.                                                #
# ----vsearch parameters 	       	       	       	       	       	       #
#    - command         Command to invoke vsearch  [default: "fastqc"].         #
#    - db_file	       Full path to a fasta file with sequences	for the	       #
#      	       	       assignation.                                            #
#    - mapFile 	       Mapping file between sequence accessions and their      #
#                      taxonomy. This file should be a two column file, tab    #
#                      separated with the accessions in the first column and   #
#                      the taxonomy in the second column with the taxonomic    # 
#                      levels separated by the 'taxo_separator', e.g.          #
#                      'AY190.44  Bacteria;Firmicutes;Bacilli;...'             #
#    - taxo_separator  Character used to split taxonomic levels	specified at   #
#      	       	       the taxonomy mapping file 'mapFile'. [default: "';'"]   #
#    - max_target_seqs Maximum number of target sequences per sequence query.  #
#      	       	       All the target sequences	are taken into account	and    #
#      	       	       their taxonomies	are mapped to their Lowest Common      #
#                      Ancestor	(LCA) with the stampa approach		       #
#		       (https://github.com/frederic-mahe/stampa).	       #
#		       "1" disables LCA mapping.      	       		       #
#    - jobs            Number of cpus. 	       	       	       	       	       #
#    - identity	       Minimum percentage of identity (between 0 and 1).       #
#    - identity_definition  Identity definitions available by vsearch. Values  #
#                      accepted are:                                           #
#                      "0". CD-HIT definition: (matching columns) / (shortest  #
#                      sequence length).                                       #
#                      "1". Edit distance: (matching columns) / (alignment     #
#                      length).                                                #
#                      "2". Edit distance excluding terminal gaps.             #
#                      "3". Marine Biological Lab definition counting each     #
#                      extended gap (internal or terminal) as a single         #
#                      difference: 1.0 - [(mismatches + gaps)/(longest sequence#
#                      length)].                                               #
#                      "4". BLAST definition, equivalent to --iddef 2 in a     #
#                      context of global pairwise alignment.                   #
#                      [default:  "2"].					       #
#    - extra_params    Any extra parameter. Run 'vsearch -h' to see all        #
#                      options [default: "--top_hits_only --maxrejects 32"]    #
# ----qiime parameters                                                         #
#    - method          Assignation method. Choose one of: "blast", "uclust" or #
#                      "rdp".                                                  #
#    - mapFile         Path to tab-delimited file mapping sequences to assigned#
#                      taxonomy. Each assigned taxonomy is provided as a       #
#                      semicolon-separated list. For assignment with "rdp",    #
#                      each assigned taxonomy must be exactly 6 levels deep.   #
#                      The default mapping file supplied by Qiime can be       #
#                      located at cascabel's environment (if created with      #
#                      conda), in such case you may found the file at:         #
#                      /../.conda/envs/cascabel/lib/python2.7/site-packages/ \ #
#                      qiime_default_reference/gg_13_8_otus/taxonomy/ \        #
#                      97_otu_taxonomy.txt.                                    #
#    - dbFile          Full path to reference fasta file or blast database     #
#                      file.                                                   #
#                      The default fasta file supplied by Qiime can be located #
#                      at cascabel's environment, if created with conda, in    #
#                      such case you may found the file at: 	               #
#                      /../.conda/envs/cascabel/lib/python2.7/site-packages/ \ #
#                      qiime_default_reference/gg_13_8_otus/taxonomy/ \        #
#                      97_otus.fasta"                                          #
#    - dbType          If 'dbFile' points to a fasta file use "-r". If 'dbFile'#
#                      points to a blast database use "-b".                    #
#    - jobs:           Number of jobs to start in parallel                     #
#------------------------------------------------------------------------------#
assignTaxonomy:
  tool: "vsearch"
  blast:
    command: "blastn"
    blast_db: ""
    fasta_db: "silva132_99.fna"
    mapFile: "taxonomy_7_levels.txt"
    taxo_separator: "';'" 
    evalue: "0.001"
    max_target_seqs: 5
    jobs: 10
    identity: 0.5   
    extra_params: ""
  vsearch:
    command: "vsearch"
    db_file: "silva132_99.fna"
    mapFile: "taxonomy_7_levels.txt"
    identity: 0.5
    taxo_separator: "';'" 
    max_target_seqs: 5 
    identity_definition: 2  
    jobs: 10
    extra_params: "--top_hits_only --maxrejects 32"
  qiime:
    method: "uclust"
    mapFile: "97_otu_taxonomy.txt."
    dbFile: "97_otus.fasta"
    dbType: "-r"
    jobs: 10
    extra_params: ""

#------------------------------------------------------------------------------#
#                               Make OTU Table                                 #
# rule: make_otu_table                                                         #
#------------------------------------------------------------------------------#
# The rule tabulates the number of times an OTU is found in each sample, and   #
# adds the taxonomic predictions for each OTU in the last column if a taxonomy #
# file is supplied.                                                            #
# - extra_params  Any extra desired parameter. Run 'make_otu_table.py --help'  #
#                 to see all the options.                                      #
#------------------------------------------------------------------------------#
makeOtu:
  extra_params: ""

#------------------------------------------------------------------------------#
#                          Filter OTUs from OTU Table                          #
# rule: filter_otu                                                             #
#------------------------------------------------------------------------------#
# This rule filters the otu table in order to remove singletons.               #
# - n             The minimum total observation count of an OTU for that otu   #
#                 to be retained [default: "2 ]                                #
# - extra_params  Any extra desired parameter. Run                             #
#                 'filter_otus_from_otu_table.py --help' to  see all options.  #
#------------------------------------------------------------------------------#
filterOtu:
  n: "2"
  extra_params: ""

#------------------------------------------------------------------------------#
#                               Biom convert                                   #
# rule: convert_table                                                          #
#------------------------------------------------------------------------------#
# This rule converts the OTU biom table into a tab separated OTU table.        #
# - tableType     ["OTU table"|"Pathway table"|"Function table"|"Ortholog      #
#                 table"|"Gene table"|"Metabolite table"|"Taxon table"|"Table"]#
#                 [default:  "--table-type 'OTU table'"].                      #
# - headerKey     The observation metadata to include from the input BIOM table#
#                 file when creating a tsv table file. [default: "--header-key # 
#                 taxonomy"].                                                  #
# - outFormat     "--to-tsv": Output as tab-separated (classic) table.         #
#                 "--to-json"  and "--to-hdf5" are also available.             #
# - extra_params: Any extra desired parameter. Run 'biom convert --help' to see#
#                 all the options                                              #
#------------------------------------------------------------------------------#
biom:
  command: "biom"
  tableType: "--table-type 'OTU table'"
  headerKey: "--header-key taxonomy"
  outFormat: "--to-tsv"
  extra_params: ""

#------------------------------------------------------------------------------#
#                           summarize_taxa.py                                  #
# rule: summarize_taxa                                                         #
#------------------------------------------------------------------------------#
# This rule provides summary information of the representation of taxonomic    #
# groups within each sample. It takes an OTU table that contains taxonomic     #
# information as input. By default, the relative abundance of each taxonomic   #
# group will be reported, but the raw counts can be returned if "-a" is passed.#
# - extra_params: Any extra desired parameter. Run 'summarize_taxa.py --help'  #
#                 to see all the options.                                      #
#                                                                              #
# The taxonomic levels to summarize are by default: "--level 2,3,4,5,6".       #
# The meaning of this level will depend on the format of the taxon strings that#
# are returned from the taxonomy assignment step. The taxonomy strings that are#
# most useful are those that standardize the taxonomic level with the depth in #
# the taxonomic strings. For instance: Level 1 = Kingdom, Level 2 = Phylum,    #
# Level 3 = Class, Level 4 = Order, Level 5 = Family, Level 6 = Genus,         #
# Level 7 = Species.                                                           #
#------------------------------------------------------------------------------#
summTaxa:
  extra_params: "--level 2,3,4,5,6,7"


#------------------------------------------------------------------------------#
#                          Filter fasta file                                   #
# rule: filter_rep_seqs                                                        #
#------------------------------------------------------------------------------#
# This rule performs OTU table-based filtering: Keep all sequences that occur  #
# in an OTU table.                                                             #
# Params:                                                                      #
# - extra_params  Any extra desired parameter. Run 'filter_fasta.py --help' to #
#                 see all options.                                             #
#------------------------------------------------------------------------------#
filterFasta:
  extra_params: ""


#------------------------------------------------------------------------------#
#                       Align representative sequences                         #
# rule: align_rep_seqs                                                         #
#------------------------------------------------------------------------------#
# This rule aligns sequences in a fasta file to each other or to a template    #
# sequence alignment, depending on the method chosen.                          #
# - align          [T/F] The user can skip this rule by setting 'align' to "F" #
#                  [default: "T"].                                             #
# -m               Method for aligning sequences. Valid choices are: "pynast", #
#                  "infernal", "clustalw", "muscle", "mafft".                  #
#                  [default: "pynast"].                                        #        
# - extra_params:  Any extra parameter. Run 'align_seqs.py --help' to see all  #
#                  options.                                                    #
#------------------------------------------------------------------------------#
alignRep:
  align: "T"
  m: "pynast"
  extra_params: ""

#------------------------------------------------------------------------------#
#                           Filter Alignment                                   #
# rule: filter_alignment                                                       #
#------------------------------------------------------------------------------#
# This rule will remove positions which are gaps in every sequence.            #
# Additionally, the user can supply a lanemask file, that defines which        #
# positions should be included when building the tree, and which should be     #
# ignored.                                                                     #
# - extra_params: Any extra desired parameter. Run 'filter_alignment.py --help'# 
#                 to see all options.                                          #
#------------------------------------------------------------------------------#
filterAlignment:
  extra_params: ""

#------------------------------------------------------------------------------#
#                           make phylogeny                                     #
# rule: make_tree                                                              #
#------------------------------------------------------------------------------#
# This rule produces a phylogenetic tree from a multiple sequence alignment.   # 
# - method:       Method for generating the tree. Valid choices are:           #
#                 "clustalw", "raxml_v730", "muscle", "fasttree", "clearcut".  # 
#                 [default: "fasttree"].                                       #
# - extra_params: Any extra parameter. Run 'make_phylogeny.py --help' to see   #
#                 all options.                                                 #
#------------------------------------------------------------------------------#
makeTree:
  method: "fasttree"
  extra_params: ""

   
  
```
