## Supplemental file 2 for "*Cascabel*: a flexible, scalable and easy-to-use amplicon sequence data analysis pipeline"

snakemake report

### Amplicon Analysis Report for Library: NIOZ66

**CASCABEL** is designed to run amplicon sequence analysis across single or multiple read libraries.

The objective of this pipeline is to create different output files which allow the user to explore data in a simple and meaningful way, as well as facilitate downstream analysis, based on the generated output files.

Another aim of **CASCABEL** is also to encourage the documentation process, by creating this report in order to assure data analysis reproducibility.

**User description:** This is the report generated by CASCABEL

Following you can see all the steps that were taken in order to get the final results of the pipeline.

#### Raw Data

The raw data for this library can be found at:

- FW raw reads: Report_NIOZ66/samples/NIOZ66/rawdata/fw.fastq

- RV raw reads: Report_NIOZ66/samples/NIOZ66/rawdata/rv.fastq

Number of total reads: 10979168.0

#### Quality Control

Evaluate quality on raw reads.

Tool: [FastQC]

Version: **FastQC v0.11.3**

**Command:**

```
fastqc Report_NIOZ66/samples/NIOZ66/rawdata/fw.fastq Report_NIOZ66/samples/NIOZ66/rawdata/rv.fastq --extract -o
Report_NIOZ66/samples/NIOZ66/qc/
```

You can follow the links below, in order to see the complete FastQC report:

- FastQC for sample NIOZ66_1: FQ1

- FastQC for sample NIOZ66_2: FQ2

**Benchmark info:**

| s | max_rss | max_vms | max_uss | max_pss | io_in | io_out | mean_load |
|---|---|---|---|---|---|---|---|
| 285.98 | 335.04 | 3310.90 | 333.00 | 333.62 | 11343.82 | 3.31 | 0.00 |

#### Read pairing

Align paired end reads and merge them into one single sequence in case they overlap.

Tool: [PEAR]

version: **PEAR v0.9.8 [April 9, 2015] - [+bzlib +zlib]**

**Command:**

```
pear -f Report_NIOZ66/samples/NIOZ66/rawdata/fw.fastq -r Report_NIOZ66/samples/NIOZ66/rawdata/rv.fastq -t 100 -v 10 -j 6 -p
0.05 -o Report_NIOZ66/runs/final_report_example/NIOZ66_data/peared/seqs >
Report_NIOZ66/runs/final_report_example/NIOZ66_data/peared/seqs.assembled.fastq
```

**Output files:**

- Merged reads: Report_NIOZ66/runs/final_report_example/NIOZ66_data/peared/seqs.assembled.fastq

- Log file: Report_NIOZ66/runs/final_report_example/NIOZ66_data/peared/pear.log

Number of peared reads: 10829329.0 = 98.64%

**Benchmark info:**

| s | max_rss | max_vms | max_uss | max_pss | io_in | io_out | mean_load |
|---|---|---|---|---|---|---|---|
| 1551.43 | 192.03 | 641.49 | 191.42 | 191.46 | 0.44 | 7177.19 | 0.00 |

#### Peared FastQC Analysis

Check the quality of the reads after assembly.

Tool: [FastQC]

Version: **FastQC v0.11.3**

**Command:**

```
fastqc Report_NIOZ66/runs/final_report_example/NIOZ66_data/peared/seqs.assembled.fastq --extract -o
Report_NIOZ66/runs/final_report_example/NIOZ66_data/peared/qc
```

**Output files:**

- FastQC report: Report_NIOZ66/runs/final_report_example/NIOZ66_data/peared/qc/seqs.assembled_fastqc.html FQ_Report

**Benchmark info:**

| s | max_rss | max_vms | max_uss | max_pss | io_in | io_out | mean_load |
|---|---|---|---|---|---|---|---|
| 186.49 | 270.09 | 3296.68 | 269.69 | 269.72 | 7068.30 | 0.68 | 0.00 |

#### Extract barcodes

Extract the barcodes used to identify individual samples.

Tool: [QIIME] - extract_barcodes.py

Version: **extract_barcodes.py 1.9.1**

**Command:**

```
extract_barcodes.py -f Report_NIOZ66/runs/final_report_example/NIOZ66_data/peared/seqs.assembled.fastq -c
barcode_single_end --bc1_len 12 -o Report_NIOZ66/runs/final_report_example/NIOZ66_data/barcodes/
```

**Output files:**

- Fastq file with barcodes: Report_NIOZ66/runs/final_report_example/NIOZ66_data/barcodes/barcodes.fastq

- Fastq file with the reads: Report_NIOZ66/runs/final_report_example/NIOZ66_data/barcodes/reads.fastq

**Benchmark info:**

| s | max_rss | max_vms | max_uss | max_pss | io_in | io_out | mean_load |
|---|---|---|---|---|---|---|---|
| 223.23 | 96.51 | 715.47 | 93.62 | 94.04 | 50.21 | 6804.02 | 0.00 |

#### Correct Barcodes

Try to correct the barcode from unassigned reads.

Maximum number of missmatches **2**.

Tool: CASCABEL's R script

**Command:**

```
Rscript Scripts/errorCorrectBarcodes.R $PWD Report_NIOZ66/metadata/sampleList_mergedBarcodes_NIOZ66.txt
Report_NIOZ66/runs/final_report_example/NIOZ66_data/barcodes/barcodes.fastq 2
```

**Output file:**

- Barcode corrected file: Report_NIOZ66/runs/final_report_example/NIOZ66_data/barcodes/barcodes.fastq_corrected

**Benchmark info:**

| s | max_rss | max_vms | max_uss | max_pss | io_in | io_out | mean_load |
|---|---|---|---|---|---|---|---|
| 2828.17 | 976.03 | 1244.32 | 968.75 | 971.26 | 868.50 | 855.34 | 0.00 |

### Demultiplexing

Library splitting, also known as demultiplexing is carried on several steps.

#### Split samples from Fastq file

Tool: [QIIME] - split_libraries_fastq.py

version: **split_libraries_fastq.py 1.9.1**

**Command:**

```
split_libraries_fastq.py -m Report_NIOZ66/metadata/sampleList_mergedBarcodes_NIOZ66.txt -i
Report_NIOZ66/runs/final_report_example/NIOZ66_data/barcodes/reads.fastq -o
Report_NIOZ66/runs/final_report_example/NIOZ66_data/splitLibs -b
Report_NIOZ66/runs/final_report_example/NIOZ66_data/barcodes/barcodes.fastq_corrected -q 19 -r 5 --retain_unassigned_reads --
barcode_type 12
```

**Benchmark info:**

| s | max_rss | max_vms | max_uss | max_pss | io_in | io_out | mean_load |
|---|---|---|---|---|---|---|---|
| 665.93 | 431.34 | 1253.81 | 425.92 | 426.96 | 6829.06 | 4248.02 | 0.00 |

#### Retain assigned reads

**Command:**

```
cat Report_NIOZ66/runs/final_report_example/NIOZ66_data/splitLibs/seqs.fna | grep -P -A1 "(?!>Unass)^>" | sed '/^--$/d' >
Report_NIOZ66/runs/final_report_example/NIOZ66_data/splitLibs/seqs.assigned.fna
```

#### Create file with only unassigned reads

**Command:**

```
cat Report_NIOZ66/runs/final_report_example/NIOZ66_data/splitLibs/seqs.fna | grep "^>Unassigned" | sed 's/>Unassigned_[0-9]*
/@/g' | sed 's/ .*//' | grep -F -w -A3 -f - Report_NIOZ66/runs/final_report_example/NIOZ66_data/peared/seqs.assembled.fastq | sed
'/^--$/d' >Report_NIOZ66/runs/final_report_example/NIOZ66_data/splitLibs/unassigned.fastq
```

#### Reverse complement unassigned reads

Tool: [Vsearch]

version: **vsearch v1.8.0_linux_x86_64, 377.8GB RAM, 36 cores**

**Command:**

```
vsearch --fastx_revcomp Report_NIOZ66/runs/final_report_example/NIOZ66_data/splitLibs/unassigned.fastq --fastqout
Report_NIOZ66/runs/final_report_example/NIOZ66_data/splitLibs/unassigned.reversed.fastq
```

#### Barcode extraction for reverse complemented, unassigned reads

Tool: [QIIME] - extract_barcodes.py

Version: **extract_barcodes.py 1.9.1**

**Command:**

```
extract_barcodes.py -f Report_NIOZ66/runs/final_report_example/NIOZ66_data/splitLibs/unassigned.reversed.fastq -c barcode_single_end --bc1_len 12 -o Report_NIOZ66/runs/final_report_example/NIOZ66_data/barcodes_unassigned/
```

#### Correct reverse complemented barcodes

Maximum number of missmatches **2**.

Tool: CASCABEL's R script

**Command:**

```
Rscript Scripts/errorCorrectBarcodes.R $PWD Report_NIOZ66/metadata/sampleList_mergedBarcodes_NIOZ66.txt Report_NIOZ66/runs/final_report_example/NIOZ66_data/barcodes_unassigned/barcodes.fastq_corrected 2
```

**Output file:**

- Barcode corrected file: Report_NIOZ66/runs/final_report_example/NIOZ66_data/barcodes/barcodes.fastq_corrected

**Benchmark info:**

| s | max_rss | max_vms | max_uss | max_pss | io_in | io_out | mean_load |
|---|---|---|---|---|---|---|---|
| 1523.63 | 897.99 | 1160.44 | 890.71 | 893.24 | 442.64 | 445.75 | 0.00 |

#### Split reverse complemented reads

Tool: [QIIME] - extract_barcodes.py

Version: **extract_barcodes.py 1.9.1**

**Command:**

```
split_libraries_fastq.py -m Report_NIOZ66/metadata/sampleList_mergedBarcodes_NIOZ66.txt -i Report_NIOZ66/runs/final_report_example/NIOZ66_data/barcodes_unassigned/reads.fastq -o Report_NIOZ66/runs/final_report_example/NIOZ66_data/splitLibsRC -b Report_NIOZ66/runs/final_report_example/NIOZ66_data/barcodes_unassigned/barcodes.fastq_corrected -q 19 -r 5 --barcode_type 12
```

**Benchmark info:**

| s | max_rss | max_vms | max_uss | max_pss | io_in | io_out | mean_load |
|---|---|---|---|---|---|---|---|
| 665.93 | 431.34 | 1253.81 | 425.92 | 426.96 | 6829.06 | 4248.02 | 0.00 |

**Output files:**

- FW reads fasta file with new header: Report_NIOZ66/runs/final_report_example/NIOZ66_data/splitLibs/seqs.assigned.fna

- Text histogram with the length of the fw reads: Report_NIOZ66/runs/final_report_example/NIOZ66_data/splitLibs/histograms.txt

- Log file for the fw reads: Report_NIOZ66/runs/final_report_example/NIOZ66_data/splitLibs/split_library_log.txt

- RV reads fasta file with new header: Report_NIOZ66/runs/final_report_example/NIOZ66_data/splitLibsRC/seqs.fna

- Text histogram with the length of the rv reads: Report_NIOZ66/runs/final_report_example/NIOZ66_data/splitLibsRC/histograms.txt

- Log file for the rv reads: Report_NIOZ66/runs/final_report_example/NIOZ66_data/splitLibsRC/split_library_log.txt

Number of reads assigned on FW: 5041715 = 46.56% of the peared reads

Number of reads assigned on RV: 4765971 = 44.01% of the peared reads

#### Generate single sample fastq files

Create single fastq files per samples (based on the raw data without applying any filtering).

Tool: CASCABEL's Java program

**Command:**

```
java -cp Scripts DemultiplexQiime --fasta -d Report_NIOZ66/runs/final_report_example/NIOZ66_data/seqs_fw_rev_accepted.fna -o Report_NIOZ66/runs/final_report_example/NIOZ66_data/demultiplexed/ -r1 Report_NIOZ66/samples/NIOZ66/rawdata/fw.fastq -r2 Report_NIOZ66/samples/NIOZ66/rawdata/fw.fastq
```

**The demultiplexed files can be located at:**

- demultiplexed directory: Report_NIOZ66/runs/final_report_example/NIOZ66_data/demultiplexed/

- summary file: Report_NIOZ66/runs/final_report_example/NIOZ66_data/demultiplexed/summary.txt

**Benchmark info:**

| s | max_rss | max_vms | max_uss | max_pss | io_in | io_out | mean_load |
|---|---|---|---|---|---|---|---|
| 1614.72 | 8318.34 | 72617.97 | 8319.68 | 8319.76 | 13.74 | 2518.16 | 0.42 |

#### Combine reads

Concatenate forward and reverse reads.

**Command:**

```
cat Report_NIOZ66/runs/final_report_example/NIOZ66_data/splitLibs/seqs.assigned.fna Report_NIOZ66/runs/final_report_example/NIOZ66_data/splitLibsRC/seqs.fna > Report_NIOZ66/runs/final_report_example/NIOZ66_data/seqs_fw_rev_accepted.fna
```

**Output file:**

- Fasta file with combined reads: Report_NIOZ66/runs/final_report_example/NIOZ66_data/seqs_fw_rev_accepted.fna

Total number of accepted reads: 9807686 = 90.57% of the peared reads or 89.33% of the raw reads

**Benchmark info:**

| s | max_rss | max_vms | max_uss | max_pss | io_in | io_out | mean_load |
|---|---|---|---|---|---|---|---|
| 6.68 | 1.63 | 12.21 | 1.12 | 1.12 | 0.00 | 4015.21 | 0.00 |

#### Remove too long and too short reads

Remove very short and long reads, with lengths more than some standard deviation below or above the mean to be short or long respectively

- Minimun length expected (shorts): 289

- Maximun length expected (longs): 319

**Command:**

```
awk '!/^>/ { next } { getline seq } length(seq) > shorts && length(seq) < longs { print $0 "n" seq }' Report_NIOZ66/runs/final_report_example/NIOZ66_data/seqs_fw_rev_accepted.fna > Report_NIOZ66/runs/final_report_example/NIOZ66_data/seqs_fw_rev_filtered.fasta
```

**Sequence distribution before remove reads**

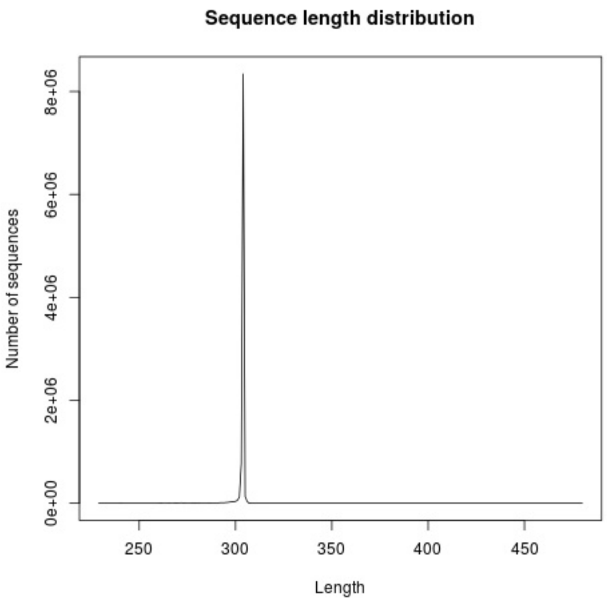


**Output file:**

- Fasta file with correct sequence length: Report_NIOZ66/runs/final_report_example/NIOZ66_data/seqs_fw_rev_filtered.fasta

Total number of reads after trimming: 9676698

Percentage of reads vs raw reads: 88.14%

Percentage of reads after demultiplexed reads vs : 98.66%

**Benchmark info:**

| s | max_rss | max_vms | max_uss | max_pss | io_in | io_out | mean_load |
|---|---|---|---|---|---|---|---|
| 17.58 | 20.43 | 123.92 | 14.84 | 16.57 | 0.00 | 3849.00 | 72.04 |

**To see a better detail on the library sample distribution please refere to the file:** Report_NIOZ66/runs/final_report_example/NIOZ66_data/seqs_fw_rev_filtered.dist.txt

#### Sample distribution

| Sample | Seqs | prc. | Sample | Seqs | prc. | Sample | Seqs | prc. | Sample | Seqs | prc. |
|---|---|---|---|---|---|---|---|---|---|---|---|
| NIOZ66.1 | 201289 | 2.08 | NIOZ66.29 | 68596 | 0.71 | NIOZ66.57 | 39726 | 0.41 | NIOZ66.85 | 41389 | 0.43 |
| NIOZ66.2 | 49588 | 0.51 | NIOZ66.30 | 153719 | 1.59 | NIOZ66.58 | 100640 | 1.04 | NIOZ66.86 | 78757 | 0.81 |
| NIOZ66.3 | 107992 | 1.12 | NIOZ66.31 | 52705 | 0.54 | NIOZ66.59 | 149962 | 1.55 | NIOZ66.87 | 33366 | 0.34 |
| NIOZ66.4 | 154077 | 1.59 | NIOZ66.32 | 25268 | 0.26 | NIOZ66.60 | 237437 | 2.45 | NIOZ66.88 | 35788 | 0.37 |
| NIOZ66.5 | 104693 | 1.08 | NIOZ66.33 | 64858 | 0.67 | NIOZ66.61 | 38222 | 0.39 | NIOZ66.89 | 96758 | 1.00 |
| NIOZ66.6 | 62500 | 0.65 | NIOZ66.34 | 118549 | 1.23 | NIOZ66.62 | 41562 | 0.43 | NIOZ66.90 | 28150 | 0.29 |
| NIOZ66.7 | 15168 | 0.16 | NIOZ66.35 | 37834 | 0.39 | NIOZ66.63 | 12179 | 0.13 | NIOZ66.91 | 12458 | 0.13 |
| NIOZ66.8 | 160212 | 1.66 | NIOZ66.36 | 48977 | 0.51 | NIOZ66.64 | 51387 | 0.53 | NIOZ66.92 | 65763 | 0.68 |
| NIOZ66.9 | 121777 | 1.26 | NIOZ66.37 | 80691 | 0.83 | NIOZ66.65 | 22147 | 0.23 | NIOZ66.93 | 99990 | 1.03 |
| NIOZ66.10 | 112564 | 1.16 | NIOZ66.38 | 85083 | 0.88 | NIOZ66.66 | 101281 | 1.05 | NIOZ66.94 | 106401 | 1.10 |
| NIOZ66.11 | 105190 | 1.09 | NIOZ66.39 | 90007 | 0.93 | NIOZ66.67 | 115423 | 1.19 | NIOZ66.95 | 92681 | 0.96 |
| NIOZ66.12 | 41491 | 0.43 | NIOZ66.40 | 134537 | 1.39 | NIOZ66.68 | 68162 | 0.70 | NIOZ66.96 | 86257 | 0.89 |

| | | | | | | | | | | | |
|---|---|---|---|---|---|---|---|---|---|---|---|
| NIOZ66.13 | 121336 | 1.25 | NIOZ66.41 | 90845 | 0.94 | NIOZ66.69 | 26572 | 0.27 | NIOZ66.97 | 120372 | 1.24 |
| NIOZ66.14 | 204831 | 2.12 | NIOZ66.42 | 32933 | 0.34 | NIOZ66.70 | 58130 | 0.60 | NIOZ66.98 | 81719 | 0.84 |
| NIOZ66.15 | 150328 | 1.55 | NIOZ66.43 | 128145 | 1.32 | NIOZ66.71 | 50495 | 0.52 | NIOZ66.99 | 47691 | 0.49 |
| NIOZ66.16 | 90884 | 0.94 | NIOZ66.44 | 168767 | 1.74 | NIOZ66.72 | 1 | 0.00 | NIOZ66.100 | 64889 | 0.67 |
| NIOZ66.17 | 153640 | 1.59 | NIOZ66.45 | 308248 | 3.19 | NIOZ66.73 | 67747 | 0.70 | NIOZ66.101 | 90468 | 0.93 |
| NIOZ66.18 | 60021 | 0.62 | NIOZ66.46 | 89463 | 0.92 | NIOZ66.74 | 38724 | 0.40 | NIOZ66.102 | 29145 | 0.30 |
| NIOZ66.19 | 116165 | 1.20 | NIOZ66.47 | 153666 | 1.59 | NIOZ66.75 | 23599 | 0.24 | NIOZ66.103 | 34595 | 0.36 |
| NIOZ66.20 | 108480 | 1.12 | NIOZ66.48 | 35159 | 0.36 | NIOZ66.76 | 157430 | 1.63 | NIOZ66.104 | 75686 | 0.78 |
| NIOZ66.21 | 97361 | 1.01 | NIOZ66.49 | 115974 | 1.20 | NIOZ66.77 | 63349 | 0.65 | NIOZ66.105 | 114011 | 1.18 |
| NIOZ66.22 | 96457 | 1.00 | NIOZ66.50 | 192098 | 1.99 | NIOZ66.78 | 26146 | 0.27 | NIOZ66.106 | 41919 | 0.43 |
| NIOZ66.23 | 48702 | 0.50 | NIOZ66.51 | 323105 | 3.34 | NIOZ66.79 | 40879 | 0.42 | NIOZ66.107 | 127248 | 1.31 |
| NIOZ66.24 | 65520 | 0.68 | NIOZ66.52 | 189804 | 1.96 | NIOZ66.80 | 58575 | 0.61 | NIOZ66.108 | 117692 | 1.22 |
| NIOZ66.25 | 39352 | 0.41 | NIOZ66.53 | 43217 | 0.45 | NIOZ66.81 | 94611 | 0.98 | NIOZ66.109 | 6991 | 0.07 |
| NIOZ66.26 | 56724 | 0.59 | NIOZ66.54 | 151219 | 1.56 | NIOZ66.82 | 30254 | 0.31 | NIOZ66.110 | 91199 | 0.94 |
| NIOZ66.27 | 60348 | 0.62 | NIOZ66.55 | 87016 | 0.90 | NIOZ66.83 | 95494 | 0.99 | NIOZ66.111 | 104652 | 1.08 |
| NIOZ66.28 | 36558 | 0.38 | NIOZ66.56 | 91198 | 0.94 | NIOZ66.84 | 35600 | 0.37 | | | |

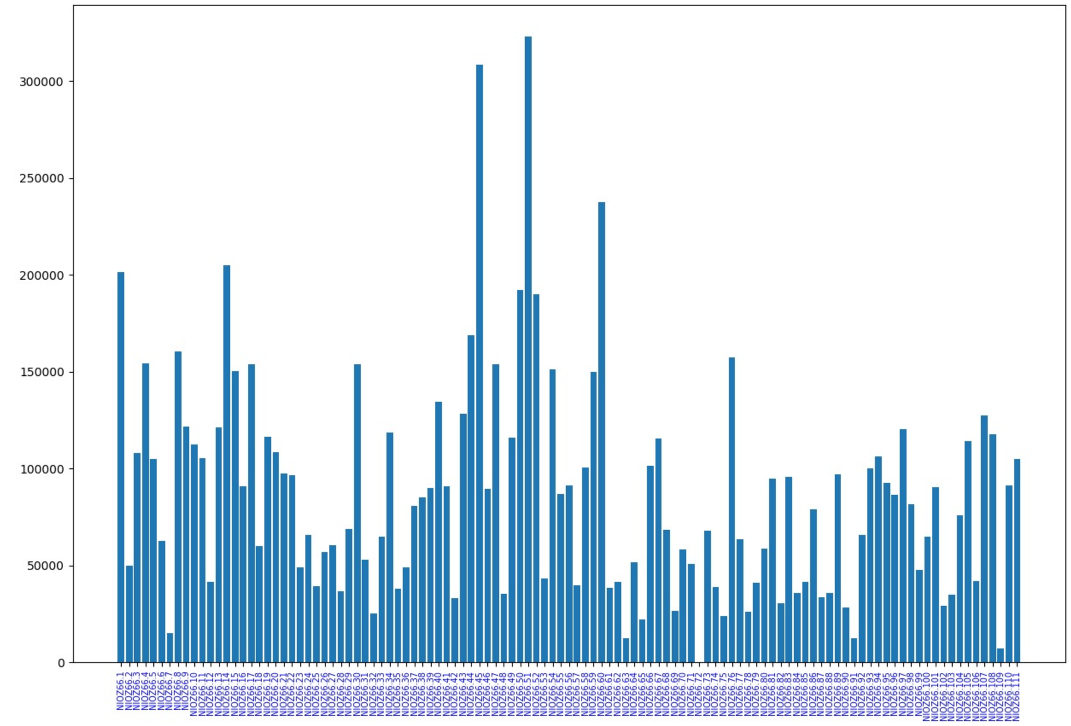


The previous chart shows the number of clean reads per sample. The bars are sorted from left to right, according to the metadata input file.

#### Final counts

Following you can see the final read counts:

| File description | Location | Number of reads | Prc(%) vs raw |
|---|---|---|---|
| Raw reads | Report_NIOZ66/samples/NIOZ66/rawdata/*.fq | 10979168.0 | 100.00% |
| Assembled reads | Report_NIOZ66/runs/final_report_example/NIOZ66_data/peared/seqs.assembled.fastq | 10829329.0 | 98.64% |
| Demultiplexed reads | Report_NIOZ66/runs/final_report_example/NIOZ66_data/seqs_fw_rev_accepted.fna | 9807686 | 89.33% |
| Length filtered | Report_NIOZ66/runs/final_report_example/NIOZ66_data/seqs_fw_rev_filtered.fasta | 9676698 | 88.14% |

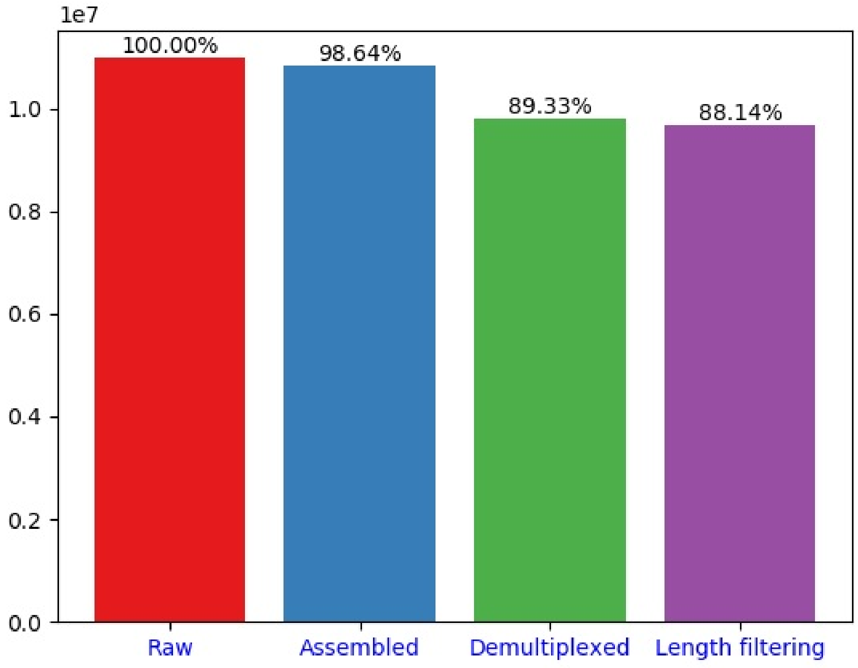


#### Taxonomic combined report

The taxonomic report performed in combination with the different libraries supplied, can be found at the following link: taxo_report (Report_NIOZ66/runs/final_report_example/report_all.html)
