## Supplemental file 3 for "*Cascabel*: a flexible, scalable and easy-to-use amplicon sequence data analysis pipeline"

snakemake report

### Amplicon Analysis Report for Libraries: final_report_example

This report consists of the OTU creation and taxonomic assignment for all the combined accepted reads of given samples or libraries, if multiple.

**User description:** This is the report generated by CASCABEL

#### Combine Reads

Merge all the reads of the individual libraries into one single file.

**Command:**

```
cat Report_NIOZ66/runs/final_report_example/NIOZ66_data/seqs_fw_rev_filtered.fasta >
Report_NIOZ66/runs/final_report_example/seqs_fw_rev_combined.fasta
```

**Output file:**

- Merged reads: Report_NIOZ66/runs/final_report_example/seqs_fw_rev_filtered.fasta

The total number of reads is: **9676698**

**Benchmark info:**

| s | max_rss | max_vms | max_uss | max_pss | io_in | io_out | mean_load |
|---|---|---|---|---|---|---|---|
| 7.12 | 18.70 | 114.36 | 13.35 | 15.07 | 0.00 | 3965.61 | 54.44 |

#### Dereplicate reads

Clusterize the reads with an identity threshold of 100%.

Tool: [vsearch]

Version: **vsearch v1.8.0_linux_x86_64, 377.8GB RAM, 36 cores**

**Command:**

```
vsearch --derep_fulllength seqs_fw_rev_combined.fasta --output seqs_fw_rev_combined_derep.fasta --uc
seqs_fw_rev_combined_derep.uc --strand both --fasta_width 0 --minuniquesize 1
```

**Output files:**

- Dereplicated fasta file: Report_NIOZ66/runs/final_report_example/derep/seqs_fw_rev_combined_derep.fasta

- Cluster file: Report_NIOZ66/runs/final_report_example/derep/seqs_fw_rev_combined_derep.uc

Total number of dereplicated sequences is: **6300129**

**Benchmark info:**

| s | max_rss | max_vms | max_uss | max_pss | io_in | io_out | mean_load |
|---|---|---|---|---|---|---|---|
| 67.01 | 3906.43 | 4201.80 | 3905.23 | 3905.32 | 0.00 | 0.00 | 0.00 |

#### Cluster OTUs

Assigns similar sequences to operational taxonomic units, or OTUs, by clustering sequences based on a user-defined similarity threshold.

Tool: [QIIME] - pick_otus.py

Version: **pick_otus.py 1.9.1**

Method: [uclust]

Identity: 0.97

**Command:**

```
pick_otus.py -m uclust -i Report_NIOZ66/runs/final_report_example/seqs_fw_rev_filtered.fasta -o
Report_NIOZ66/samples/final_report_example/otu/ -s 0.97
```

**Output files:**

- OTU List: Report_NIOZ66/runs/final_report_example/otu/seqs_fw_rev_filtered_otus.txt

- Log file: Report_NIOZ66/runs/final_report_example/otu/seqs_fw_rev_filtered_otus.log

The total number of different OTUS is: **2510664**

**Benchmark info:**

| s | max_rss | max_vms | max_uss | max_pss | io_in | io_out | mean_load |
|---|---|---|---|---|---|---|---|
| 61178.27 | 13334.91 | 15059.80 | 13332.24 | 13332.58 | 721.61 | 4669.54 | 99.48 |

#### Pick representatives

Pick a single representative sequence for each OTU.

Tool: [QIIME] - pick_rep_set.py

Version: **pick_rep_set.py 1.9.1**

Method: longest

**Command:**

```
pick_rep_set.py -m longest -i Report_NIOZ66/runs/final_report_example/otu/seqs_fw_rev_filtered_otus.txt -f
Report_NIOZ66/samples/final_report_example/seqs_fw_rev_filtered.fasta -o
Report_NIOZ66/samples/final_report_example/otu/representative_seq_set.fasta --log_fp
Report_NIOZ66/samples/final_report_example/otu/representative_seq_set.log
```

**Output file:**

- Fasta file with representative sequences: Report_NIOZ66/runs/final_report_example/otu/representative_seq_set.fasta

**Benchmark info:**

| s | max_rss | max_vms | max_uss | max_pss | io_in | io_out | mean_load |
|---|---|---|---|---|---|---|---|
| 106.48 | 5619.50 | 6238.64 | 5617.26 | 5617.59 | 8.84 | 0.02 | 0.00 |

#### Assign taxonomy

Given a set of sequences, assign the taxonomy of each sequence.

Tool: [vsearch]

Version: **vsearch v1.8.0_linux_x86_64, 377.8GB RAM, 36 cores**

Reference fasta file: /export/data01/databases/silva/qiime/SILVA_132_QIIME_release/rep_set/rep_set_all/99/silva132_99.fna

Taxonomy mapping file: /export/data01/databases/silva/qiime/SILVA_132_QIIME_release/taxonomy/taxonomy_all/99/taxonomy_7_levels.txt

**Command:**

```
vsearch--usearch_global Report_NIOZ66/runs/final_report_example/otu/representative_seq_set.fasta --db
/export/data01/databases/silva/qiime/SILVA_132_QIIME_release/rep_set/rep_set_all/99/silva132_99.fna --dbmask none --qmask
none --rowlen 0 --id 0.5 --iddef 2 --userfields query+id2+target --maxaccepts 5 --threads 10 --top_hits_only --maxrejects 32 --
output_no_hits --userout representative_seq_set_tax_vsearch.out
```

After vsearch assignation, **results were mapped to their LCA using stampa_merge.py** script

The percentage of successfully assigned OTUs is: **100.00%**

**Output file:**

- OTU taxonomy assignation:

Report_NIOZ66/runs/final_report_example/otu/taxonomy_vsearch/representative_seq_set_tax_assignments.txt

**Benchmark info:**

| s | max_rss | max_vms | max_uss | max_pss | io_in | io_out | mean_load |
|---|---|---|---|---|---|---|---|
| 4063.62 | 2246.57 | 2983.93 | 2245.87 | 2245.95 | 594.53 | 258.00 | 0.00 |

#### Make OTU table

Tabulates the number of times an OTU is found in each sample, and adds the taxonomic predictions for each OTU in the last column.

Tool: [QIIME] - make_otu_table.py

Version: **make_otu_table.py 1.9.1**

**Command:**

```
make_otu_table.py -i Report_NIOZ66/runs/final_report_example/otu/taxonomy_vsearch/seqs_fw_rev_filtered_otus.txt -t
Report_NIOZ66/runs/final_report_example/otu/taxonomy_vsearch/representative_seq_set_tax_assignments.txt -o
Report_NIOZ66/runs/final_report_example/otu/taxonomy_vsearch/otuTable.biom
```

**Output file:**

- Biom format table: Report_NIOZ66/runs/final_report_example/otu/taxonomy_vsearch/otuTable.biom

**Benchmark info:**

| s | max_rss | max_vms | max_uss | max_pss | io_in | io_out | mean_load |
|---|---|---|---|---|---|---|---|
| 86.88 | 5941.17 | 6560.16 | 5938.88 | 5939.21 | 2.24 | 579.47 | 0.00 |

#### Convert OTU table

Convert from the BIOM table format to a human readable format.

Tool: [BIOM]

Version: **biom, version 2.1.5**

**Command:**

```
biom convert -i Report_NIOZ66/runs/final_report_example/otu/taxonomy_vsearch/otuTable.biom -o
Report_NIOZ66/runs/final_report_example/otu/taxonomy_vsearch/otuTable.txt --table-type 'OTU table' --header-key taxonomy --to-
tsv
```

**Output file:**

- TSV format table: Report_NIOZ66/runs/final_report_example/otu/taxonomy_vsearch/otuTable.txt

**Benchmark info:**

| s | max_rss | max_vms | max_uss | max_pss | io_in | io_out | mean_load |
|---|---|---|---|---|---|---|---|
| 572.14 | 17859.48 | 18829.80 | 17857.15 | 17857.50 | 0.00 | 0.00 | 0.00 |

#### Summarize Taxa

Summarize information of the representation of taxonomic groups within each sample.

Tool: [QIIME] - summarize_taxa.py

Version: **summarize_taxa.py 1.9.1**

**Command:**

```
summarize_taxa.py -i Report_NIOZ66/runs/final_report_example/otu/taxonomy_vsearch/otuTable.biom --level 2,3,4,5,6,7 -o
Report_NIOZ66/runs/final_report_example/otu/taxonomy_vsearch/summary/
```

**Output file:**

- Taxonomy summarized counts at different taxonomy levels: Report_NIOZ66/runs/final_report_example/otu/taxonomy_vsearch/summary/otuTable_L**N**.txt

Where **N** is the taxonomy level. Default configuration produces levels from 2 to 6.

**Benchmark info:**

| s | max_rss | max_vms | max_uss | max_pss | io_in | io_out | mean_load |
|---|---|---|---|---|---|---|---|
| 2833.63 | 9841.44 | 10462.72 | 9838.51 | 9839.01 | 0.21 | 6.33 | 0.00 |

#### Filter OTU table

Filter OTUs from an OTU table based on their observed counts or identifier.

Tool: [QIIME] - filter_otus_from_otu_table.py

Version: **filter_otus_from_otu_table.py 1.9.1**

Minimum observation counts: 2

**Command:**

```
filter_otus_from_otu_table.py -i Report_NIOZ66/runs/final_report_example/otu/taxonomy_vsearch/otuTable.biom -o
Report_NIOZ66/runs/final_report_example/otu/taxonomy_vsearch/otuTable_noSingletons.biom -n 2
```

**Output file:**

- Biom table: Report_NIOZ66/runs/final_report_example/otu/taxonomy_vsearch/otuTable_noSingletons.biom

**Benchmark info:**

| s | max_rss | max_vms | max_uss | max_pss | io_in | io_out | mean_load |
|---|---|---|---|---|---|---|---|
| 148.09 | 7357.32 | 7975.28 | 7355.01 | 7355.35 | 0.02 | 19.32 | 0.00 |

#### Convert Filtered OTU table

Convert the filtered OTU table from the BIOM table format to a human readable format

Tool: [BIOM]

Version: **biom, version 2.1.5**

**Command:**

```
biom convert -i Report_NIOZ66/runs/final_report_example/otu/taxonomy_vsearch/otuTable_noSingletons.biom -o
Report_NIOZ66/runs/final_report_example/otu/taxonomy_vsearch/otuTable_noSingletons.txt --table-type 'OTU table' --header-key
taxonomy --to-tsv
```

**Output file:**

- TSV format table: Report_NIOZ66/runs/final_report_example/otu/taxonomy_vsearch/otuTable_noSingletons.txt

**Benchmark info:**

| s | max_rss | max_vms | max_uss | max_pss | io_in | io_out | mean_load |
|---|---|---|---|---|---|---|---|
| 148.09 | 7357.32 | 7975.28 | 7355.01 | 7355.35 | 0.02 | 19.32 | 0.00 |

#### Filter representative sequences

Remove sequences according to the filtered OTU biom table.

Tool: [QIIME] - filter_fasta.py

Version: **filter_fasta.py 1.9.1**

**Command:**

```
filter_fasta.py -f Report_NIOZ66/samples/final_report_example/otu/representative_seq_set.fasta -o
Report_NIOZ66/samples/final_report_example/otu/taxonomy_vsearch/representative_seq_set_noSingletons.fasta -b
Report_NIOZ66/samples/final_report_example/otu/otuTable_noSingletons.biom
```

**Output file:**

- Filtered fasta file: Report_NIOZ66/samples/final_report_example/otu/taxonomy_vsearch/representative_seq_set_noSingletons.fasta

**Benchmark info:**

| s | max_rss | max_vms | max_uss | max_pss | io_in | io_out | mean_load |
|---|---|---|---|---|---|---|---|
| 22.30 | 1297.94 | 1917.56 | 1295.98 | 1296.31 | 0.01 | 96.02 | 0.00 |

#### Align representative sequences

Align the sequences in a FASTA file to each other or to a template sequence alignment.

Tool: [QIIME] - align_seqs.py

Version: TBD

Method: [pynast]

**Command:**

```
align_seqs.py -m pynast -i Report_NIOZ66/runs/final_report_example/otu/vsearch/representative_seq_set_noSingletons.fasta -o
Report_NIOZ66/runs/final_report_example/otu/taxonomy_vsearch/aligned/representative_seq_set_noSingletons_aligned.fasta
```

**Output files:**

- Aligned fasta file: Report_NIOZ66/runs/final_report_example/otu/taxonomy_vsearch/aligned/representative_seq_set_noSingletons_aligned.fasta

- Log file: Report_NIOZ66/runs/final_report_example/otu/taxonomy_vsearch/aligned/representative_seq_set_noSingletons_log.txt

**Benchmark info:**

| s | max_rss | max_vms | max_uss | max_pss | io_in | io_out | mean_load |
|---|---|---|---|---|---|---|---|
| 10830.32 | 11307.14 | 13418.08 | 11301.81 | 11302.89 | 37.74 | 1717.93 | 9.48 |

#### Filter alignment

Removes positions which are gaps in every sequence.

Tool: [QIIME] - filter_alignment.py

Version: **filter_alignment.py 1.9.1**

**Command:**

```
filter_alignment.py -i
Report_NIOZ66/runs/final_report_example/otu/taxonomy_vsearch/aligned/representative_seq_set_noSingletons_aligned.fasta -o
Report_NIOZ66/runs/final_report_example/otu/taxonomy_vsearch/aligned/filtered/
```

**Output file:**

- Aligned fasta file: Report_NIOZ66/runs/final_report_example/otu/taxonomy_vsearch/aligned/representative_seq_set_noSingletons_aligned_pfiltered.fasta

**Benchmark info:**

| s | max_rss | max_vms | max_uss | max_pss | io_in | io_out | mean_load |
|---|---|---|---|---|---|---|---|
| 2092.65 | 5072.89 | 5692.19 | 5070.06 | 5070.52 | 0.49 | 0.02 | 0.00 |

#### Make tree

Create phylogenetic tree (newick format).

Tool: [QIIME] - make_phylogeny.py

Version: **make_phylogeny.py 1.9.1**

Method: [fasttree]

**Command:**

```
make_phylogeny.py -i
Report_NIOZ66/runs/final_report_example/otu/taxonomy_vsearch/aligned/representative_seq_set_noSingletons_aligned.fasta -o
representative_seq_set_noSingletons_aligned_pfiltered.tre -t fasttree
```

**Output file:**

- Taxonomy tree: Report_NIOZ66/runs/final_report_example/otu/taxonomy_vsearch/aligned/representative_seq_set_noSingletons_aligned.tre

**Benchmark info:**

| s | max_rss | max_vms | max_uss | max_pss | io_in | io_out | mean_load |
|---|---|---|---|---|---|---|---|
| 22835.48 | 5778.29 | 6412.04 | 5775.00 | 5775.43 | 0.23 | 238.39 | 99.50 |

#### Krona report

Krona allows hierarchical data to be explored with zooming, multi-layered pie charts.

Tool: [Krona]

These charts were created using the OTU table **without** singletons

The report was executed for all the samples.

Each sample is represented on a separated chart (same html report).

You can see the report at the following link:

- Krona report: kreport

Or access the html file at:

- Krona html file: Report_NIOZ66/runs/final_report_example/otu/taxonomy_vsearch/krona_report.html

**Benchmark info:**

| s | max_rss | max_vms | max_uss | max_pss | io_in | io_out | mean_load |
|---|---|---|---|---|---|---|---|
| 140.15 | 59.36 | 192.19 | 52.72 | 54.97 | 1.88 | 239.11 | 10.24 |

#### Final counts

Following the read counts:

| File description | Location | # | (%) |
|---|---|---|---|
| Combined clean reads | Report_NIOZ66/runs/final_report_example/seqs_fw_rev_combined.fasta | 9676698 | 100% |
| Dereplicated reads | Report_NIOZ66/runs/final_report_example/derep/seqs_fw_rev_combined_derep.fasta | 6300129 | 65.11% |
| OTU table | Report_NIOZ66/runs/final_report_example/otu/seqs_fw_rev_combined_remapped_otus.txt | 2510664 | 25.95% |
| Taxonomy assignation | Report_NIOZ66/runs/final_report_example/otu/taxonomy_vsearch/representative_seq_set_tax_assignments.txt | 2510662 | 100.00% |

| | | | |
|---|---|---|---|
| OTU table (no singletons: a > 2) | Report_NIOZ66/runs/final_report_example/otu/taxonomy_vsearch/otuTable_noSingletons.txt | 506239 | 20.16% |
| Assigned no singletons | Report_NIOZ66/runs/final_report_example/otu/taxonomy_vsearch/otuTable_noSingletons.txt | 506239 | 100.00% |

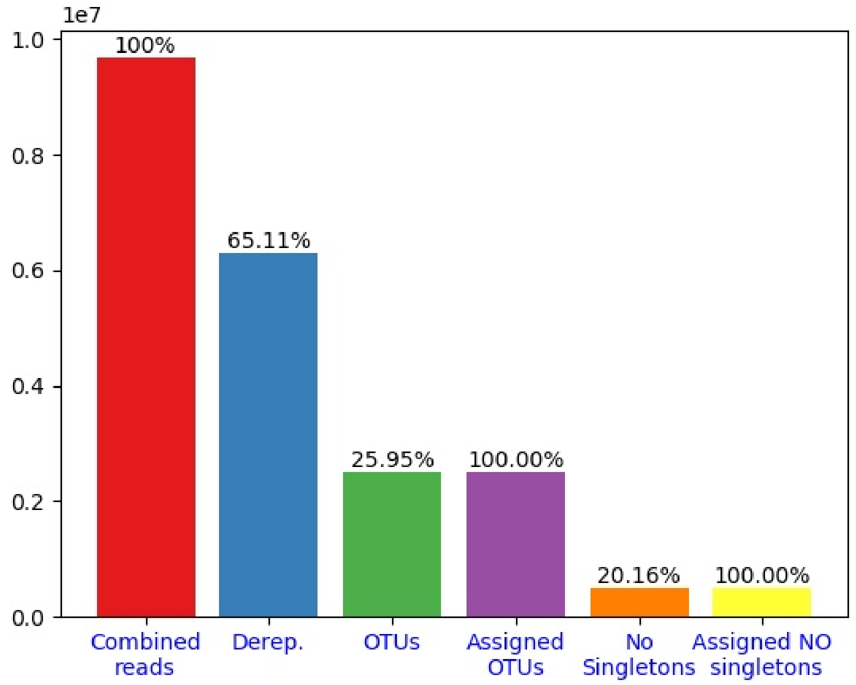


Note:

- Assigned OTUs percentage is the amount of successfully assigned OTUs.

- No singletons percentage is the percentage of no singletons OTUs in reference to the complete OTU table.

- Assigned No singletons is the amount of successfully no singletons assigned OTUs.
